# Loss of heterozygosity exposes germline mutations in complex I and drives Warburg metabolism in oncocytic carcinoma of the thyroid

**DOI:** 10.64898/2025.12.06.692631

**Authors:** Celia de la Calle Arregui, Anderson R. Frank, Kelvin Mun, Jiwoong Kim, Kuntal Majmudar, Justin A. Bishop, Cheryl Lewis, Yang Xie, David G. McFadden

**Affiliations:** Department of Internal Medicine, Division of Endocrinology, University of Texas Southwestern Medical Center, Dallas, TX 75390, USA; Department of Biochemistry, University of Texas Southwestern Medical Center, Dallas, TX 75390, USA; Section of Molecular Medicine, Department of Radiation Oncology, University of Texas Southwestern Medical Center, Dallas, TX 75390, USA; Harold C. Simmons Comprehensive Cancer Center, University of Texas Southwestern Medical Center, Dallas, TX 75390, USA; Department of Pathology, University of Texas Southwestern Medical Center, Dallas, TX 75390, USA; Quantitative Biomedical Research Center, University of Texas Southwestern Medical Center, Dallas, TX 75390; Department of Health Data Science and Biostatistics, O’Donnell School of Public Health, University of Texas Southwestern Medical Center, Dallas, TX 75390, USA

## Abstract

Oncocytic (Hürthle cell) carcinoma of the thyroid (OCT) is characterized by widespread loss of heterozygosity (LOH), mitochondrial accumulation and recurrent mitochondrial DNA mutations leading to impairment of complex I. Here, we establish and characterize a novel OCT cell line, UT946, which displays severe mitochondrial electron transport chain dysfunction and a Warburg metabolic phenotype. Using a series of cytoplasmic hybrids, we establish that the complex I defect in UT946 stems from a nuclear-encoded loss of function mutation in the complex I subunit NDUFS1. To our surprise, the mutation in NDUFS1 was inherited as a recessive germline allele that underwent LOH in the tumor to expose functional loss of complex I. A re-analysis of 91 OCT tumor genomes revealed that LOH-driven exposure of recessive germline mutations in complex I subunits was a recurrent mechanism underlying complex I inactivation in OCT. These findings unveil a new germline-driven mechanism of complex I loss and metabolic reprogramming in cancer, and provide further evidence of the strong selective pressure for complex I impairment in OCT.

**Teaser:** Germline mutations in complex I induce aerobic glycolysis in oncocytic carcinoma of the thyroid through somatic loss of heterozygosity.

## Introduction

Central carbon metabolism, which includes glycolysis, the tricarboxylic acid (TCA) cycle, and the mitochondrial electron transport chain, is an interconnected series of biochemical reactions that allows cells to extract energy from nutrients such as glucose and produce the intermediate metabolites required for growth and proliferation^1^. Defects in central carbon metabolism typically have profound effects on an organism’s growth and development. For example, mutations in components of the mitochondrial ETC can cause disorders including Leigh syndrome and mitochondrial encephalopathy, lactic acidosis, and stroke-like episodes (MELAS), which present as systemic disorders affecting multiple organ systems including the central nervous and musculoskeletal systems^2^.

Early studies by Otto Warburg revealed that tumors produce excessive amounts of lactate even under aerobic conditions, a phenomenon later termed the “Warburg effect,” which he attributed to impaired mitochondrial respiration^3^. Cellular respiration is performed by the mitochondrial ETC, a series of multiprotein complexes (NADH dehydrogenase, succinate dehydrogenase, cytochrome *bc_1_* oxidoreductase, cytochrome *c* oxidase, and ATP synthase; also referred to as complexes I-V, respectively) that transfer electrons from reduced carriers such as NADH + H^+^ and FADH_2_ to oxygen. Ultimately, ATP production is coupled to these electron transfer steps, allowing cells to efficiently generate energy from carbon-based fuels^1^. While subsequent studies have confirmed that tumors and cultured cancer cells take up glucose and produce lactate at higher rates than healthy tissues, there is significant evidence demonstrating that mitochondrial ETC activity is required for tuHmor growth and metastasis^4–11^. Consistent with this, loss-of-function (LOF) mutations in mitochondrial ETC complex subunits, which are encoded by both the nuclear and mitochondrial genomes, are relatively uncommon in tumors^12–14^.

However, specific mutations in TCA cycle and mitochondrial ETC components are pro-tumorigenic, including gain-of-function mutations in isocitrate dehydrogenase 1 and 2 (*IDH1* and *IDH2*) and loss-of-function mutations in succinate dehydrogenase/mitochondrial ETC complex II (*SDHA, SDHB, SDHC, and SDHD)* or fumarate hydratase (*FH*)^15^. The consequences of these mutations, which include the production of the oncogenic metabolite, (R)-2-hydroxyglutarate ((R)-2-HG) by mutant IDH, and the accumulation of the TCA cycle intermediates succinate and fumarate resulting from impaired SDH or FH activity, contribute to tumor development and progression through competitive inhibition of alpha-ketoglutarate (α-KG)-dependent enzymes such as dioxygenases and demethylases that ultimately regulate gene expression^15–18^. These mutations are typically restricted to tumors arising from specific tissues, suggesting that the cell-of-origin may determine sensitivity to transformation by the metabolic consequences and gene expression programs induced by (R)-2-HG-dependent modulation of enzymes.

Our group and others previously reported that deleterious mitochondrial DNA (mtDNA) mutations affecting mitochondrial ETC complex I (NADH dehydrogenase) were enriched in OCT and maintained during tumor progression and metastasis^19–21^. This contrasts with multiple other tumor types from diverse lineages that do not display enrichment of damaging mtDNA mutations, despite the fact that somatic mtDNA mutations in tumors are relatively common^12,14,22^. Oncocytomas, a class of tumors which includes OCT, are characterized by extensive mitochondrial accumulation, and also display enrichment of deleterious mtDNA mutations^19–21^. In addition to having an abundance of mitochondria, OCT tumors are glucose-avid and display pronounced ^18^F-FDG uptake on PET scans^23–25^. These observations raise the possibility that genetic impairment of the mitochondrial ETC could underlie the mitochondrial accumulation and a metabolic reprogramming observed in OCT. Despite these common features, damaging somatic mtDNA mutations have been observed in approximately 60% of OCT tumors^19,20^, suggesting that 40% of OCT tumors either do not exhibit mitochondrial ETC impairment, or any defects are driven by alternative genetic or epigenetic mechanisms. Here, we report the generation of a novel OCT patient-derived cell line that exhibits impaired complex I function. Our efforts to uncover the genetic basis of CI impairment in this model led to the discovery of a novel mechanism of CI loss in OCT.

## Results

### UT946 exhibits mitochondrial ETC dysfunction

We established a cell line derived from a primary OCT tumor with poorly differentiated/anaplastic features, UT946. Because previous studies have reported severe mitochondrial ETC dysfunction in patient-derived OCT models^26–28^ we examined mitochondrial ETC function and cellular metabolism in a panel of thyroid cancer cell lines including UT946 (OCT), NCI-237^UTSW^ (OCT), TPC-1 (papillary thyroid carcinoma/PTC) and UT354 (poorly-differentiated thyroid carcinoma (PDTC). We performed Seahorse extracellular flux assays^29,30^ and observed that UT946, along with the OCT cell line NCI-237^UTSW^, exhibited a severe respiration defect and increased extracellular acidification rate relative to non-OCT cell lines (Figure 1A and B). Cells with mitochondrial ETC defects require exogenous pyruvate and glucose for proliferation. Pyruvate sustains NAD^+^ regeneration and ultimately, aspartate biosynthesis^31–34^, whereas glucose supports cellular ATP production via glycolysis^35–37^. We tested whether UT946 required exogenous pyruvate and glucose for viability and growth by culturing cells in pyruvate-free or glucose-free media. Both OCT cell lines including UT946 required exogenous pyruvate for proliferation (Figure 1C). Additionally, we observed that UT946, as well as NCI-237^UTSW^, exhibited a significant reduction in viability when grown in glucose-free media supplemented with galactose (Figure 1D). By contrast, ETC-proficient thyroid cancer cell lines remained viable in media without glucose and sustained proliferation in pyruvate-free media (Figure 1C and D).

**Figure 1.**
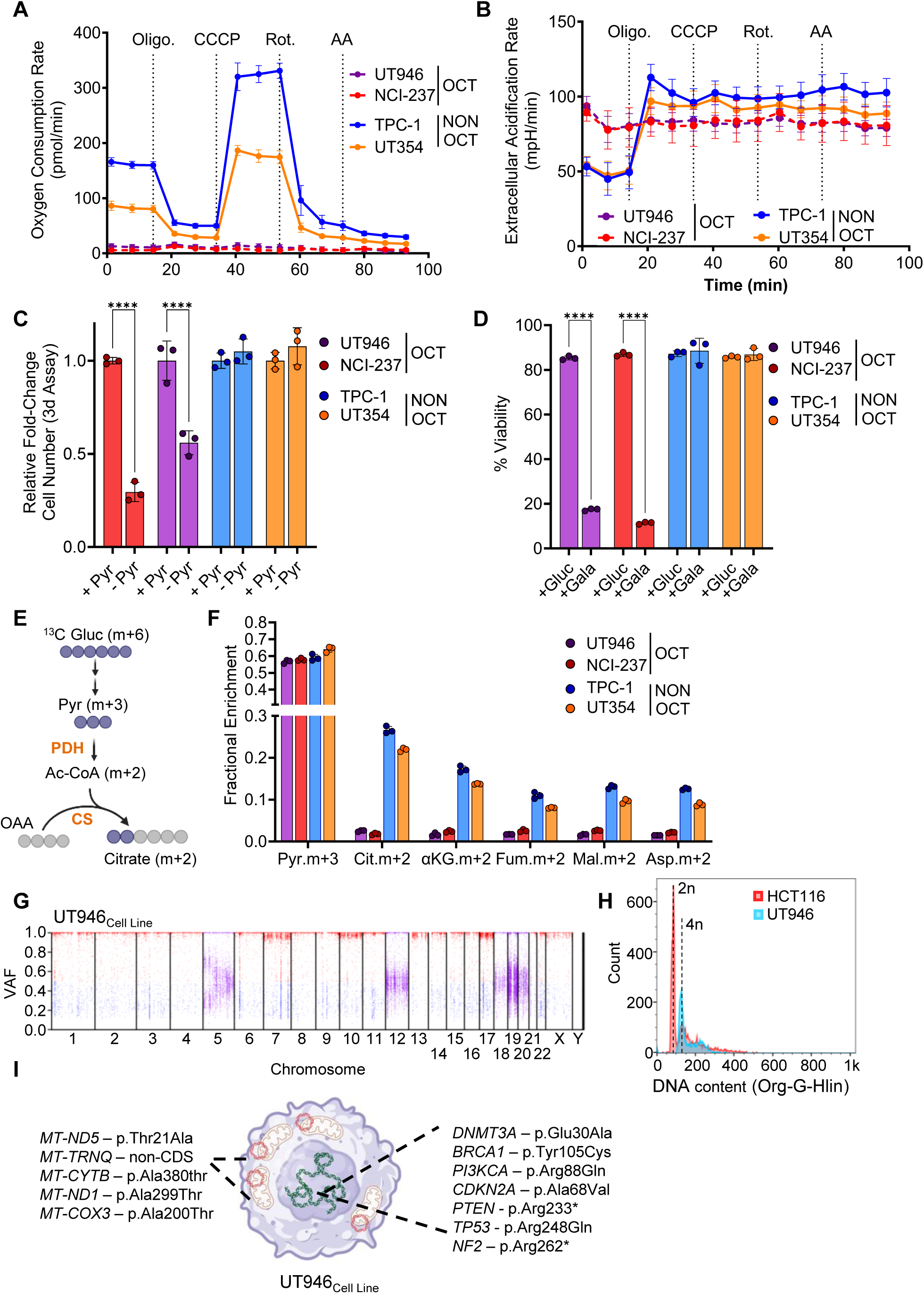
UT946 exhibits ETC dysfunction and genetic features of OCT. **A-B,** Oxygen consumption rate and acidification rate in intact cells after the addition of Oligo = oligomycin; CCCP = carbonyl cyanide 3-chlorophenylhydrazone; Rot. = Rotenone and Anti A = antimycin A (HPLM) n=6-8. **C,** Fold-change in cell number of indicated cell lines in media ± 100 μmol/L pyruvate. n = 3 (HPLM). **D,** Cell viability of indicated cell lines after 48h in media ± 5mM glucose or 5mM galactose n=3 (HPLM). **E,** Schematic representation of ^13^C glucose labelled carbons into glycolysis and TCA cycle. **F,** Mass isotopomer abundance for the indicated metabolites in cells cultured with [U- ^13^C_6_] glucose for 6 hours (HPLM). Pyr. = Pyruvate, Cit. = Citrate, αKG = α-ketoglutarate, Fum. = Fumarate, Mal. = malate, Asp. = Aspartate. **G,** Loss of heterozygosity plot showing variant allele fraction (VAF) of germline variants for UT946 Cell Line. Variants with VAF >0.5 are in red; variants with VAF <0.5 are in blue. **H,** DNA content measured by propidium iodine staining in a diploid (HCT116) cell vs UT946 cell line. **I,** Schematic representation of selected somatic mutations found in UT946 Cell Line. Data are plotted as mean ± SD of the indicated number of replicates

We also assessed glucose metabolism in the panel of cell lines by culturing cells in media containing ^13^C-labeled glucose ([U-^13^C_6_] glucose) and analyzing glucose carbon fate using gas chromatography-mass spectrometry (GC-MS). UT946 and NCI-237^UTSW^ displayed minimal incorporation of glucose-derived carbon into the citrate pool via pyruvate dehydrogenase (PDH) and citrate synthase (CS) (Figure 1E and F) indicating overall reduced glucose-derived carbon movement through the TCA cycle. Altogether, we conclude that UT946 exhibited impaired mitochondrial ETC function and alterations in glucose metabolism consistent with a defect in complex I.

### UT946 harbors key genetic features of OCT

We hypothesized that UT946 harbored a mtDNA-encoded CI defect, like 60% of OCT tumors and all OCT cells lines reported to date^19,20^. To establish a nuclear and mitochondrial genomic profile we performed whole-exome sequencing (WES) of the primary tumor and matched cell line, referred to as UT946_Tumor_ and UT946_Cell Line_, respectively. Normal thyroid tissue was not collected at the time of tumor resection. We analyzed exome sequencing datasets for regions of homozygosity and we determined that UT946_Tumor_ displayed loss-of-heterozygosity (LOH) across Chr 1-4, 6, 8, 9, 11, 14-16, and 20-22, consistent with widespread LOH previously reported for larger cohorts of OCT tumors (Supplementary Figure 1A)^19,20,26^. UT946_Cell Line_ displayed more extensive LOH, with additional loss of one copy of Chr 7, 10, 13, 17, and X, while retaining heterozygosity for Chr 20 (Figure 1G). To our knowledge, UT946 represents the only OCT reported to date that exhibits extensive LOH on Chr 7. While many OCT tumors develop near-haploid genomes as a result of widespread chromosome loss, OCT tumors frequently undergo whole-genome duplication (WGD) following chromosome loss, resulting in tumors with LOH and complex ploidies^19,20^. We analyzed the cellular DNA content of UT946_Cell Line_ using propidium iodide staining and flow cytometry and observed approximately 4N DNA content, suggestive of UT946_Cell Line_ having undergone WGD (Figure 1H).

Because normal thyroid or blood were not collected from this patient, we identified putative somatic mutations from WES data by extensive annotation of known germline single nucleotide polymorphisms (SNPs) to exclude common germline mutations(Supplementary Table 1). Missense mutations of unknown functional significance from known single nucleotide polymorphisms (SNP) in the DNA methyltransferase *DNMT3A*, (Chr2:25300227T>G; p.Glu30Ala; VAF^WES^ = 0.990), as well as in the DNA repair gene *BRCA1* (Chr17:43104249T>C; p.Tyr105Cys; VAF^WES^ = 0.466) (Supplementary Figure 1B) were found in UT946_tumor_. These mutations were maintained in UT946_Cell Line_. Consistent with LOH on Chr 17 in UT946_Cell Line_, the VAF of the *BRCA1* mutation was ∼1. Potentially deleterious missense or nonsense mutations in a series of tumor suppressors including *CDKN2A* (Chr9:21971156G>A; p.Ala68Val; VAF^WES^ = 0.565), *PTEN* (Chr10:87957915C>T; p.Arg233*; VAF^WES^ = 0.960), *TP53* (Chr17:7674220C>T; p.Arg248Gln; VAF^WES^ = 0.950), and *NF2* (Chr22:29661313C>T; p.Arg262*; VAF^WES^ = 0.992) were identified in UT946_Cell Line_, as well as a non-canonical activating mutation in *PIK3CA* that creates an arginine-to-glutamine substitution at amino acid residue 88 (Chr3:179199088G>A; p.Arg88Gln; VAF^WES^ = 0.451) (Figure 1I and Supplementary Figure 1B)^38–40^. These mutations have previously been reported in advance forms of thyroid carcinomas, consistent with the pathological observations of poorly differentiated thyroid carcinoma (PDTC) and anaplastic thyroid carcinoma (ATC) features^41,42^. Together with the observation of additional regions of LOH in UT946_Cell Line_, the emergence of these mutations associated with advanced forms of thyroid cancer is consistent with the outgrowth of a tumor sub-clone during the generation of the cell line.

We next sought to identify mitochondrial DNA mutations present in the UT946 tumor and cell line. Off-target exome sequencing reads that mapped to mtDNA exhibited a mean coverage depth of 1,212X and 123X in UT946_Tumor_ and UT946_Cell Line_, respectively. Analysis of mtDNA-mapped reads identified two variants of interest in UT946_Tumor_: a homoplasmic missense mutation in *MT-ND5* causing a threonine-to-alanine substitution at amino acid residue 21 (ChrM:12397A>G; p.Thr21Ala; VAF^WES^ = 0.999) and a non-coding homoplasmic mutation in *MT-TRNQ* (ChrM:4336T>C; VAF^WES^ = 1) (Figure 1I and Supplementary Figure 1C). While both mutations were flagged and listed in ClinVar as having possible associations with Leigh’s syndrome and mitochondrial encephalopathy, lactic acidosis, and stroke-like episodes (MELAS), respectively, there were limited data to support these associations. These mutations were maintained at homoplasmy during establishment of UT946_Cell Line_, and we identified three additional heteroplasmic mtDNA mutations including missense mutations in *MT-ND1* (ChrM:4201G>A; p.Ala299Thr; VAF^WES^ = 0.232), *MT-COX3* (ChrM:9804G>A; p.Ala200Thr; VAF^WES^ = 0.180), and *MT-CYTB* (ChrM:15884G>A; p.Ala380Thr; VAF^WES^ = 0.432) (Supplementary Figure 1C). We concluded from our metabolic and genomic analysis that UT946 represented a bona fide OCT tumor: cell line pair.

### Mitochondrial ETC dysfunction in UT946 is not mtDNA-encoded

We hypothesized that the MT-ND5^T21A^ missense mutation represented a LOF mutation resulting in complex I impairment in UT946. To test whether this mutation was causal for the mitochondrial ETC dysfunction in UT946, we generated cytoplasmic hybrid (cybrid) cell lines that harbored different mitochondrial genomes in a common nuclear background^7,43^. Cybrids were prepared by fusion of mtDNA-depleted host cells (ρ^0^ cells) and enucleated mitochondrial donor cells (cytoplasts) followed by fluorescence activated cell sorting (FACS) of putative cybrids and subsequent selection in uridine-free media^7^. This strategy allowed us to isolate cybrid lines with different combinations of nuclear and mitochondrial genomes (Figure 2A). To ensure the results were not biased due to cell line subclones, we generated two or three independent cybrid cell lines for each mitochondrial:nuclear genome pair of interest. UT946, TPC-1, or 143B (an osteosarcoma cell line regularly utilized in cybrid experiments) enucleated cytoplasts were obtained by ultracentrifugation against a Percoll gradient. ρ^0^ cells were obtained by serial passaging of UT946 and TPC-1 cells in 2’,3’-dideoxycytidine (ddC), a nucleoside analog that inhibits mtDNA replication^44,45^. To ensure mtDNA depletion in ρ^0^ cells, we measured mtDNA levels by qPCR using primer pairs directed against mtDNA targets (*MT-ND1* and *MT-CO3*) and a nuclear genome target (*ACTB*). ddC treatment caused progressive depletion of mtDNA up to 500-fold relative to untreated cells. Removal of ddC led to a slow but consistent recovery of the mtDNA over time (Supplementary Figure 2A and B). After cybrid generation and expansion, we confirmed mtDNA repopulation in cybrid lines by qPCR. UT946-based cybrids displayed nearly 100% mtDNA repopulation whereas TPC-1-based cybrids were not fully repopulated (Supplementary Figure 2C). Importantly, TPC-1-based cybrids displayed mitochondrial ETC function similar to parental TPC-1 cells (discussed below), suggesting that the level of mtDNA repopulation was sufficient to restore mitochondrial function. To validate the nuclear and mitochondrial genomic integrity of the cybrids, we used short-tandem repeat (STR) profiling and Oxford nanopore long-read sequencing of PRC-amplified mtDNA fragments respectively (Supplementary Table 2 and Supplementary Figure 2D and E). We discarded cybrids containing incorrect or mixed nucleus and/or mitochondrial DNAs, and we successfully established several cybrid lines to investigate ETC dysfunction in UT946_Cell Line_ (Figure 2A).

**Figure 2.**
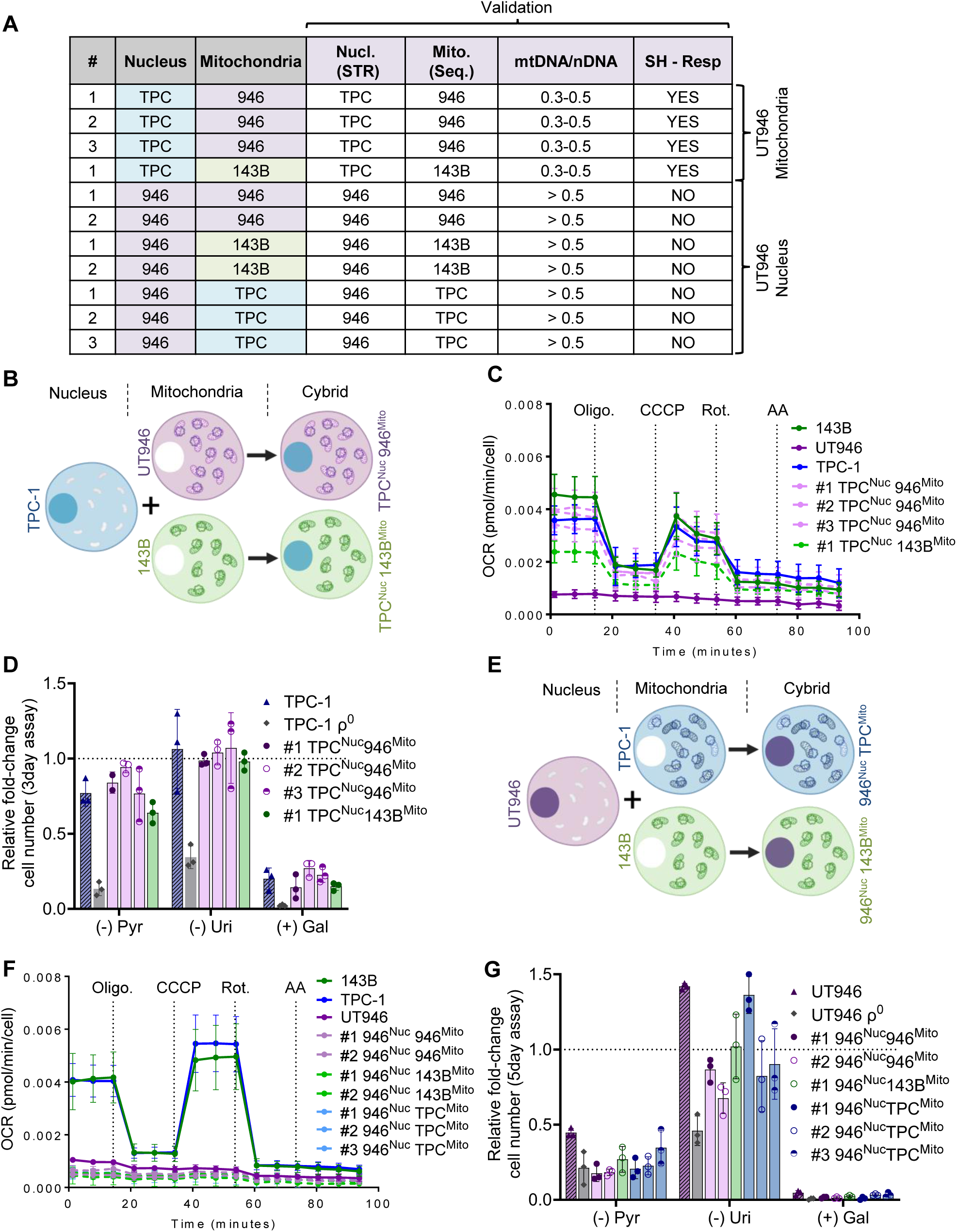
Mitochondrial ETC dysfunction in UT946 is not mtDNA-encoded. **A,** Summary table of generated cybrids and the subsequent technical validation. **B,** Schematic representation of TPC cybrids **C,** Normalized oxygen consumption rate in indicated cybrids and cell lines n=8-12. **D,** Fold change in cell number relative to complete media of indicated cell lines in media containing ± 100 μmol/L pyruvate, 50 µg/mL Uridine, 5 mM glucose/galactose n = 3. **E,** Schematic representation of UT946 cybrids **F,** Normalized oxygen consumption rate in indicated cybrids and cell lines n=8-12. **G,** Fold change in cell number relative to complete media of indicated cell lines in media containing ± 100 μmol/L pyruvate, 50 µg/mL Uridine, 5 mM glucose/galactose n = 3. Oligo = oligomycin; CCCP = carbonyl cyanide 3-chlorophenylhydrazone; Rot. = Rotenone, Anti A = antimycin A. #CB = # Cybrid. Data are plotted as mean ± SD of the indicated number of replicates

To determine whether UT946 mtDNA encoded a functional ETC, we introduced UT946 mitochondria into TPC-1 nuclear donors (TPC^Nuc^946^mito^). If the MT-ND5^T21A^, or any other variant, encoded in the UT946 mtDNA acted as a loss-of-function event, TPC^Nuc^946^mito^ cybrids should exhibit a respiration defect. In contrast, we observed that TPC-1-based cybrids harboring UT946 mtDNA displayed oxygen consumption and extracellular acidification rates similar to the parental TPC-1 cell line or TPC-1 cybrids expressing functional 143B mtDNA (TPC^Nuc^143B^mito^) (Figure 2B, C and Supplementary Figure 2F). Consistent with intact mitochondrial ETC function, we also observed that TPC^Nuc^946^mito^ cybrid lines exhibited no growth defect, relative to parental TPC-1 cells or a TPC^Nuc^143B^mito^ cybrid line, when cultured in pyruvate- or uridine- free media, or in galactose-containing media (Figure 2D). We concluded that, contrary to our hypothesis, the UT946 mitochondrial genome encoded a fully functional complex I.

### Mitochondrial complex I dysfunction in UT946 is nuclear encoded

We next tested whether the UT946 nuclear genome encoded a defect in mitochondrial respirational ETC. We introduced functional mitochondria from 143B and TPC-1 cells into UT946 nuclear donors, generating UT946-based cybrids (946^nuc^143B^mito^ and 946^nuc^TPC^mito^). These UT946-based cybrids exhibited oxygen consumption defects and extracellular acidification rates which closely matched the parental UT946 cell line or UT946 ρ^0^ cells repopulated with UT946 mitochondria (946^nuc^946^mito^) (Figure 2E, F and Supplementary Figure 2G). All UT946-based cybrids displayed proliferation defects when cultured in pyruvate-free or galactose-containing media, but not uridine-free media, suggesting a mitochondrial ETC defect linked to complex I (Figure 2G). Together, these findings indicated that the ETC defect in UT946 cells originated from a nuclear-encoded defect in complex I rather than a damaging mtDNA mutation.

To identify potential genetic causes of complex I dysfunction in UT946_Cell Line_, we re-examined the WES data and focused on nuclear-encoded complex I subunits. We identified a missense mutation in the complex I subunit, *NDUFS1*, (Chr2:206146957A>G; p.Val228Ala) that generated a valine-to-alanine substitution at position 228. In both the cell line and tumor sequencing datasets, this variant existed at an allele fraction of 1.0, suggesting that the Chr2 LOH we observed (Figure 1G) led to the loss of the wild-type allele. Although the chemical change between valine and alanine was modest, previous studies had identified NDUFS1^V228A^ as a disease-associated mutation in compound heterozygous patients with complex I deficiency^46–48^. In addition, computational predictions CADD^49^ (28.1), REVEL^50^ (0.944), and PolyPhen-2^51^ (0.999) indicated a high likelihood of pathogenicity for this variant.

To examine the effect of NDUFS1^V228A^ mutation on complex I function, we disrupted *NDUFS1* using CRISPR-Cas9 in respiration-competent TPC-1 cells. Cells with NDUFS1 deletion displayed significantly reduced oxygen consumption relative to wild-type TPC-1 cells or TPC-1 cells in which we targeted the neutral AAVS1 locus (Figure 3A-C), confirming the essentiality of NDUFS1 gene for CI function. We next expressed an exogenous copy of either wild-type NDUFS1 or NDUFS1^V228A^ in NDUFS1 KO cells (Figure 3A). Re-expression of wild-type NDUFS1, but not NDUFS1^V228A^, was sufficient to restore respiration and rescue growth in pyruvate-free or glucose-free (galactose-replete) replete media in TPC-1 NDUFS1 KO cells (Figure 3B-D, and supplementary Figure 3A). A proliferation defect was not observed upon uridine withdrawal, suggesting the ETC defect induced by NDUFS1^V228A^ was linked specifically to complex I (Figure 3D). These results established that NDUFS1^V228A^ acted as a loss of function event. However, whether this mutation was solely responsible for the CI defect observed in UT946 cells was not certain.

**Figure 3.**
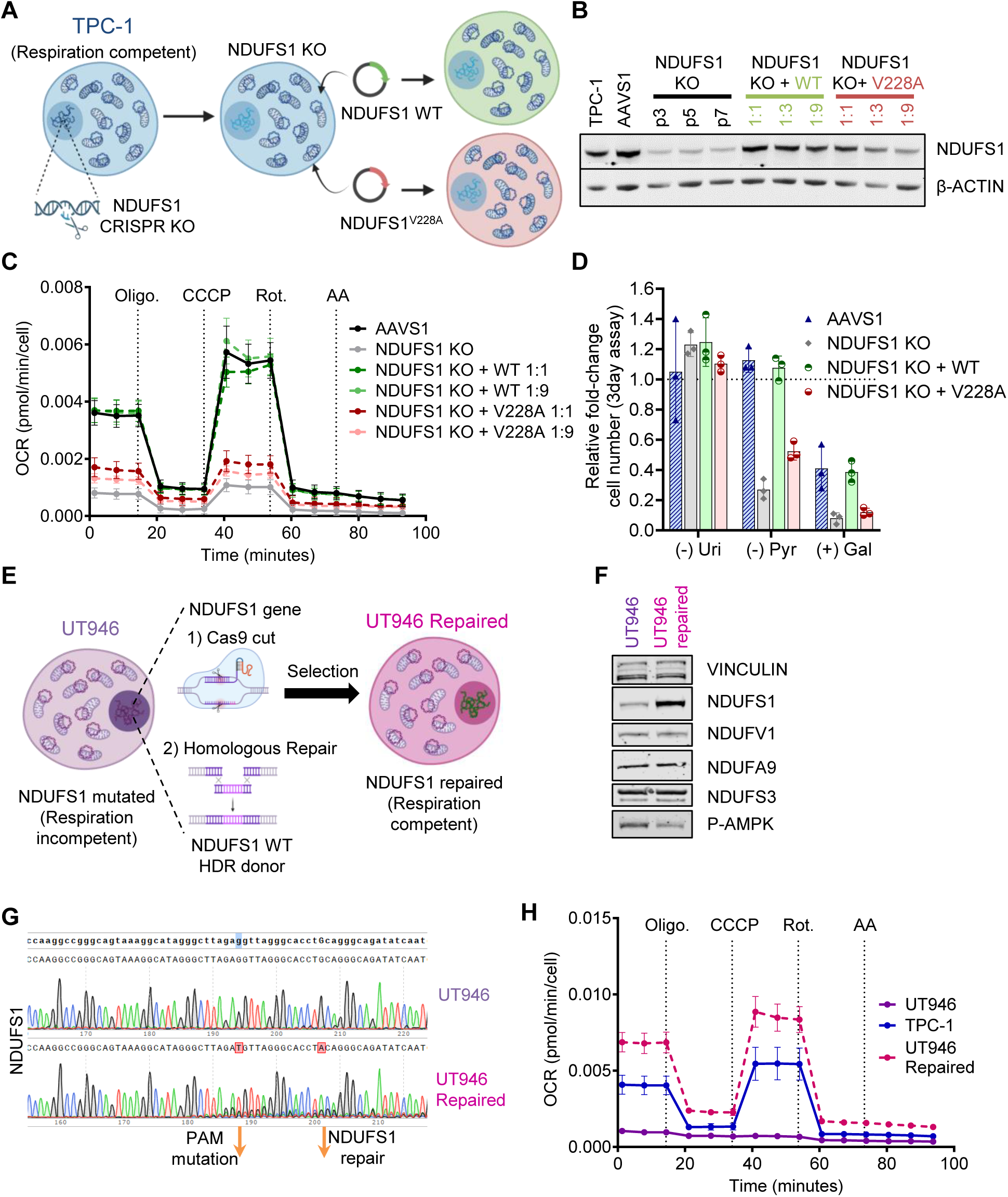
Mitochondrial complex I dysfunction in UT946 is nuclear-encoded. **A,** Schematic representation of NDUFS1^V228A^ validation strategy. NDUFS1 gene was knocked out (KO) in TPC-1 respiration competent cells followed by the e-expression of the wild type (WT) or V228A form. **B,** TPC-1 edited cells were collected at indicated passages (p3, p5, p7) after transduction with CRISPR-Cas9 system. KO cells were infected as indicated viral dilution (1:1, 1:6, 1:9) with either the WT or V228A form of NDUFS1. Protein lysates were immunoblotted for indicated proteins. **C,** Normalized oxygen consumption rate in indicated TPC-1-edited cell lines 1:1, 1:9 refers to viral dilution upon NDUFS1 re-expression n=8-12. **D,** Fold change in cell number relative to complete media of indicated cell lines in media containing ± 100 μmol/L pyruvate (Pyr.), 50 µg/mL Uridine (Uri.), 5 mM glucose or 5mM galactose (Gal.) n = 3. **E,** Schematic representation of the repair strategy. HDR = Homologous-directed repair. **F,** UT946 repaired cells were collected after selection. Protein lysates were immunoblotted for indicated proteins**. G,** Sanger sequencing traces of the genomic region surrounding the V228A mutation in the indicated cell lines. PAM = protospacer adjacent motif. **H,** Normalized oxygen consumption rate in indicated UT946-edited cell lines n=8-12. Oligo = oligomycin; CCCP = carbonyl cyanide 3-chlorophenylhydrazone; Rot. = Rotenone, Anti A = antimycin A. Data are plotted as mean ± SD of the indicated number of replicates

We therefore used CRISPR-Cas9 to revert the NDUFS1^V228A^ mutation. We introduced Cas9 ribonucleoprotein (RNP) complex targeting *NDUFS1* with a single-stranded oligodeoxynucleotide (ssODN) template encoding the wild-type NDUFS1 sequence at amino acid 228 (Figure 3E) into UT946 cells. Cells harboring wild-type NDUFS1 were enriched via a metabolism-based selection strategy wherein cells were cultured in glucose-free (galactose-replete) media. If reversion of the NDUFS1^V228A^ mutation to the WT allele led to restoration of CI function, these cells should survive in glucose-free media whereas cells with mutant NDUFS1 should die. We observed significant, but not complete cell death after UT946 repaired cells were shifted to glucose-free conditions. Surviving cells were expanded and pooled. We observed increased NDUFS1 protein levels in UT946 repaired cells compared to UT946 cells harboring the NDUFS1^V228A^ mutation (Figure 3F). This finding raised the possibility that a complex I defect in UT946 cells resulted either from destabilization of the NDUFS1 protein or indirectly through destabilization of the entire complex as has been previously reported following mutation of individual subunits^52^. We assessed NDUFS1 mutation status in UT946 repaired cells by Sanger sequencing of PCR amplicons surrounding NDUFS1 amino acid 228. Indeed, we observed reversion of the V228A mutation to the WT allele (Figure 3G). We subjected UT946 control and UT946 repaired cells to Seahorse extracellular flux assays to determine whether repair of the mutation was sufficient to restore CI and ETC function. Indeed, we observed restored respiratory activity in UT946 cells in which NDUFS1 was repaired via CRISPR-Cas9 (Figure 3H and Supplementary Figure 3B). Also consistent with an intact ETC, UT946 NDUFS1^repair^ cells were no longer auxotrophic for glucose or pyruvate (Supplementary Figure 3C). These results established that NDUFS1^V228A^ was singularly responsible for the CI defect observed in UT946 cells, and demonstrated that at least in some cases, nuclear mutations in CI subunits can drive metabolic alterations in OCT.

### Loss of heterozygosity of recessive germline mutations unmasks CI deficiency in OCT

Complex I is a multi-subunit enzyme composed of 45 proteins, of which 38 are encoded by the nuclear genome and seven by the mitochondrial genome. Although sequencing studies of OCT tumors have consistently identified CI mutations in approximately 60–70% of cases, C1 mutations have only been reported in mitochondrial-encoded CI genes ^19,20^. We hypothesized that the UT946 tumor represented a rare example of a somatic mutation in a nuclear-encoded Complex I subunit. However, because matched normal thyroid tissue was initially unavailable, we could not establish somatic origin of the NDUFS1^V228A^ mutation. We retrieved paraffin-embedded tumor and adjacent normal thyroid tissue from the clinical tissue blocks. We performed targeted PCR amplification and Sanger sequencing of *NDUFS1* from normal thyroid and tumor to determine if the NDUFS1^V228A^ mutation arose specifically in the tumor. We observed NDUFS1^V228A^ in normal thyroid as a heterozygous mutation which rejected our hypothesis that NDUFS1^V228A^ was acquired during tumorigenesis (Figure 4A). Although we could not discriminate between a germline variant and a developmental field mutation, we favored the hypothesis that NDUFS1^V228A^ was an inherited variant. NDUFS1 is encoded on chromosome 2, which underwent whole chromosome loss in the UT946 primary tumor and cell line. This LOH event resulted in the loss of the wild-type NDUFS1 allele, exposing NDUFS1^V228A^ and CI impairment.

**Figure 4.**
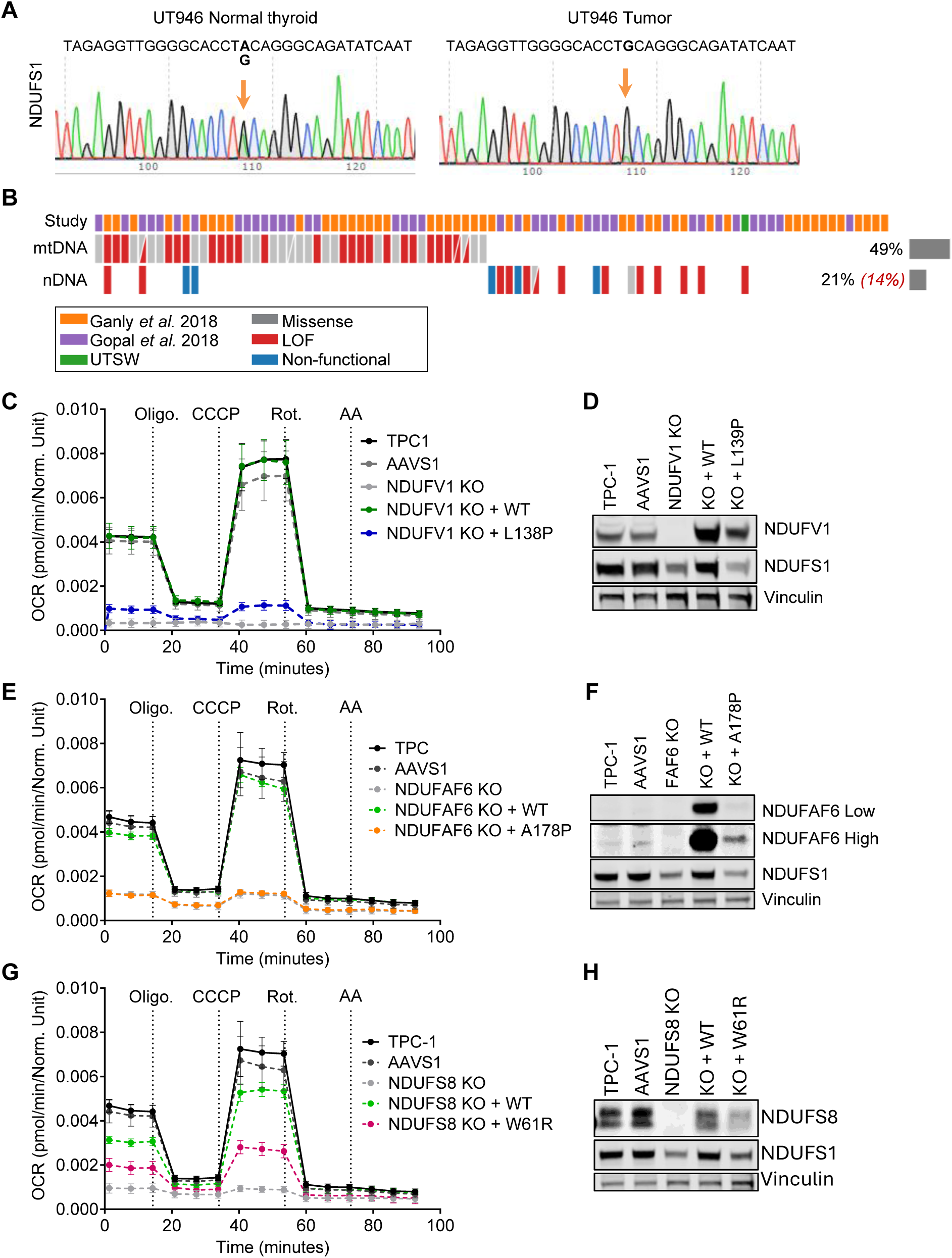
Loss of heterozygosity of recessive germline mutations unmasks CI deficiency. **A,** Sanger sequencing traces of NDUFS1^V228A^ mutation in UT946 normal thyroid and tumor regions. DNA obtained from paraffin blocks. **B,** CoMut plot showing somatic mitochondrial encoded mutations in Complex I genes (mtDNA) and recessive germline variants that underwent LOH in the tumor targeting Complex I genes encoded in the nucleus (nDNA). Silent (validated) mutations are missense mutations validated to be non-functional. **C,** Normalized oxygen consumption rate in NDUFV1 TPC-1-edited cell lines n=8-12. **D,** Protein lysates form NDUFV1 TPC-1 edited cells were immunoblotted for indicated proteins. **E,** Normalized oxygen consumption rate in NDUFAF6 TPC-1-edited cell lines n=8-12. **F,** Protein lysates form NDUFAF6 TPC-1 edited cells were immunoblotted for indicated proteins. **G,** Normalized oxygen consumption rate in NDUFS8 TPC-1-edited cell lines n=8-12. **H,** Protein lysates form NDUFS8 TPC-1 edited cells were immunoblotted for indicated proteins. Oligo = oligomycin; CCCP = carbonyl cyanide 3-chlorophenylhydrazone; Rot. = Rotenone, Anti A = antimycin A. Data are plotted as mean ± SD of the indicated number of replicates

The observation of a recessive, loss-of-function allele (NDUFS1^V228A^) in tumor-adjacent normal thyroid prompted us to explore whether LOH driven exposure of recessive germline variants represented an alternative mechanism of complex I impairment in OCT. Tumor genome data are typically analyzed to identify true somatic mutations; consequently, variants present in matched normal tissue or buffy coat samples are excluded. We reanalyzed whole-exome sequencing (WES) data from previously published studies of OCT, comprising a total of 91 OCT patients^19,20^ (Figure 4B and Supplementary Table 3). We specifically annotated germline variants that underwent LOH in our analysis. We observed a total of 19/91 (21%) cases that harbored germline coding variants in CI subunits that underwent loss of the WT allele in the tumor samples. Of these, 8/91 (9%) represented clear LOF mutations, including four frameshift mutations and one nonsense mutation in subunits required for complex I function: NDUFV2 (p.His21Argfs*6), NDUFA6 (p.Met104Cysfs*35), NDUFAF7 (p.Gln384Alafs*7), NDUFAF3 (p.Gly164Serfs*29) and NDUFB6 (p.Tyr84*)^53–56^ (Supplementary Figure 4A). Additionally, we identified a non-start splicing-altering mutation in NDUFAF6 (chr8:95046993 G>A) and 2 start codon mutations in NUBPL and NDUFAF4^57–61^. In at least three of these cases, buffy coat was used at the normal tissue control which established a germline origin for these events (Supplementary Figure 4A).

We assessed whether mutations in nuclear-encoded complex I genes were mutually exclusive with mtDNA encoded mutations, which would support a similar selective advantage of these events. We re-analyzed exome sequencing reads that mapped to mtDNA to identify somatic mtDNA events. To identify mutations that were likely to impact CI function, we selected missense and LOF mutations in protein-coding sequences with variant allelic fraction (VAF) >0.7 in tumors and VAF <0.5 in matched normal (Figure 4B and Supplementary Table 3). Although our initial pipeline failed to call 10 variants reported in the original studies, after individual review, we confirmed and included each of these events in our analysis Figure 4B. Of the 19 individuals with nuclear-encoded CI mutations, 15 (80%) had either no detectable mtDNA mutations or low-heteroplasmy missense variants, which were unlikely to have functional consequences. Among the four patients harboring mutations in both genomes, only two (HCTC-102 and SRS3086539) carried loss-of-function mutations in both nuclear- and mitochondrial-encoded CI genes (Figure 4B and Supplementary Table 3).

We observed 11/91 (12%) cases that harbored a germline CI missense mutation that underwent LOH in tumor samples. Notably, two independent patients harbored the same NDUFS1^V228A^ mutation we identified and studied in UT946. In one case, NDUFS1^V228A^ was detected as a heterozygous mutation in the buffy coat, confirming germline origin (Supplementary Figure 4A). The NDUFS1^V228A^ mutation was therefore detected in 3.26% (3/91) of OCT patients, representing a ∼250-fold enrichment, compared with a reported frequence of 0.013% in the general population (www.gnomAD.org).

We also identified 8 additional missense mutations of unknown significance affecting CI core, supernumerary, and assembly factors (Supplementary Figure 4A). NDUFV1 (p.Leu138Pro), NDUFS8 (p.Trp61Arg), NDUFV3 (p.Pro4Arg, p.Arg26Gln), NDUFA6 (p.His108Tyr), NDUFAF6 (p.Ala178Pro), NUBPL (p.Asn198Thr) and TMEM126B (p.Asp133Asn). Each of these subunits, except NDUFV3, was reported to be required for CI function^52,56,57,59,62–64^. The necessity of these subunits for CI function raised the possibility that some of these variants induced a functional CI defect in OCT tumors. To determine if these mutations encoded a CI defect, we used CRISPR-Cas9 to disrupt each of the genes in respiration-competent TPC-1 cells and subsequently re-express the corresponding wild-type or mutant subunit.

Multiple *in silico* tools supported the deleterious nature of NDUFV1^L138P^ and TMEM126B^D133N^ variants, including high scores from CADD (31.0 and 29.2), REVEL (0.936 and 0.861) and PolyPhen-2 (0.991 and 1), respectively. Re-expression of NDUFV1^L138P^ failed to restore respiration in NDUFV1 KO cells, consistent with L138P representing a loss of function variant (Figure 4C and D and Supplementary Figure 4B-E). In contrast, TPC1-TMEM126B^KO^ cells expressing TMEM126B^D133N^ exhibited similar respiratory activity as cells expressing the wild-type protein, suggesting that D133N was a silent mutation. NDUFAF6^A178P^ was predicted to be a deleterious mutation; however, computational evidence was less compelling compared to variants such as NDUFV1^L138P^ CADD (26.2), REVEL (0.73) and PolyPhen-2 (0.979). Nevertheless, re-expression of NDUFAF6^A178P^ in NDUFAF6 KO cells failed to rescue respiration in NDUFAF6 KO cells (Figure 4E and F and Supplementary Figure 4F) establishing this variant as a *bona fide* loss-of-function mutation. NDUFA6^H108T^ and NUBPL^N198T^ variants were not predicted to be pathogenic. Consistently, re-expression of the mutant alleles in the corresponding KO cells restored respiration (Supplementary Figure 4G-N). Finally, the NDUFS8^W61R^ variant was not present in gnomAD or clinVar databases. We observed that upon re-expression of the mutant form, respiration in NDUFS8 KO cells was partially restored, suggesting that, within the limit of our experimental strategy to precisely control expression levels, NDUFS8^W61R^ might act as a hypomorphic variant (Figure 4G and H and Supplementary Figure 4O). In all cases, deletion of the target CI subunit resulted in destabilization of other CI subunits, including NDUFS1, as has been previously reported. Consequently, NDUFS1 protein levels served as a surrogate marker for NDUFAF6 KO in the absence of a high-quality NDUFAF6-specific antibody (Supplementary Figure 4H). As anticipated, not every missense mutation led to functional impairment of CI. Nonetheless, 13 of 91 (14%) OCT tumors carried definitive loss-of-function mutations in nuclear-encoded CI genes, with one additional tumor harboring a hypomorphic mutation. Together, these findings suggest that ETC dysfunction in OCT generally arises through one of two distinct mechanisms: (1) somatically acquired mutations in mtDNA-encoded CI subunits, or (2) LOH-driven exposure of germline loss-of-function mutations in nuclear-encoded CI subunits.

## Discussion

Here, we report the generation of an OCT tumor cell line that exhibits the key features of these uncommon cancers: impaired mitochondrial respiration and widespread loss of chromosomes^19–21^. The generation of UT946 represents an advance for OCT research, considering that only three patient-derived models have been reported^26,27,65^. Our efforts to determine the genetic basis of CI impairment in UT946 fortuitously led to the discovery of a recessive LOF germline allele in *NDUFS1* that underwent LOH in the tumor to expose the mutant phenotype. We put forth that this study provides an example of the power of studying rare disease outliers with extreme phenotypes (and genotypes). By probing the genetic basis of CI impairment in this tumor-cell line pair derived from a single patient, we uncovered an unexpected genetic mechanism of somatic CI impairment leading to the Warburg effect. This mechanism, however, is not uncommon in OCT, and represents a recurrent genetic mechanism that explains the basis of CI impairment and Warburg metabolism in 14% (13/92) of our OCT cohorts. Together with the approximately 60% of cases that harbor putative LOF mutations in the mtDNA, we now understand the genetic basis of complex I loss in approximately 75% of cases. Whether the remaining 25% harbor mutations in proteins not yet known to control complex I function, or in other pathways that lead to increased mitochondrial biogenesis that is pathognomonic of OCT will require additional studies. Testing for germline events that undergo LOH might be warranted in clinical genotyping assays should therapies be developed that target CI, as has been reported in laboratory models^26,27^. The UT946 isogenic pair in which repair of NDUFS1^V228A^ fully restored CI function might serve as an isogenic experimental system to uncover new CI-specific dependencies.

LOH that exposes germline mutations represents a classical mechanism of tumor suppressor inactivation. Mutations in complex II (succinate dehydrogenase complex) underlie familial paraganglioma syndromes^66^. However, SDH also acts as a TCA cycle enzyme, and tumor development following SDH loss is reported to stem from an accumulation of the SDH substrate succinate to levels that impair ⍺-ketoglutarate (⍺-KG) dependent enzymes^15^. This mechanism is similar to mutations in other TCA enzymes including fumarate hydratase (FH) and isocitrate dehydrogenase (IDH) in which the enzyme substrates accumulate and impair ⍺-KG dependent dioxygenases and other enzymes. CI loss, which does not lead to accumulation of TCA substrates, induces distinct metabolic alterations compared to mutations within TCA cycle enzymes^67^. These differences, and the absence of recurrent SDH, FH, or IDH mutations in OCT suggest CI loss imparts different selective advantages compared to defects in TCA cycle enzymes.

Our finding of recurrent LOF germline mutations in CI that undergo LOH to expose the mutant allele provides further evidence of the selective advantage of CI loss in OCT. This observation parallels our previous report of allelic enrichment of CI mutations encoded in the mtDNA in OCT, in comparison to most cancers in which CI loss is selected against^12,26^. We also note that the NDUFS1^V228A^ mutation was identified in three independent patients in our cohort (∼3%). Considering www.gnomAD.org reports an allele frequency of 1.3 x 10^-4^ for this variant, we observe over 250-fold enrichment for this allele in our small cohort of patients. Positive selection for CI mutations has also been demonstrated in experimental models^68,69^. However, experimental evidence demonstrating that CI loss provides a growth advantage is limited, suggesting that CI loss could be under selection in very early stages of tumorigenesis, or that CI loss becomes fixed in the tumor at a very early stage of tumorigenesis. Importantly, OCT does not classically present in a familial pattern, and in the absence of a family in which CI germline mutations are strongly associated with tumor incidence, we specifically avoid claiming that these mutations represent bona fide tumor suppressor events akin to SDH mutations. We favor that inheritance of a recessive CI LOF allele might predispose to the development of oncocytic features in a thyroid tumor, and in this sense represents a tumor modifying allele as opposed to a tumor suppressor mutation. Nonetheless, additional studies of patients with recessive CI alleles will be required to determine whether these mutations predispose to thyroid nodules, benign or malignant, and if oncocytic features are more common in nodules that arise in these patients.

Why haven’t these germline mutations in CI been previously described? Cancer genome sequencing studies generally seek to identify tumor-specific somatic mutations. Therefore, germline events are excluded in analysis pipelines to identify somatic mutations. Despite the fact that a majority of the OCT genome undergoes LOH, all prior studies, including our own, followed a standard analysis approach and failed to report germline events^19,20^. In addition, identification of recurrent mutations in genes that encode subunits of large multiprotein complexes is more complex and requires detailed knowledge of subunit and complex assembly, stability and function.

This study exemplifies that power that the germline retains over tumor genotype and phenotype, and reminds us to review germline variation in tumor genome studies, particularly in tumors, like OCT, in which a large fraction of the genome undergoes LOH^19,20^. We hypothesize that continued analysis of tumor genomes to systematically identify LOH of germline alleles will uncover additional genes and pathways that contribute to tumorigenesis and modify tumor phenotypes.

## Materials and Methods

### Patient-Derived Materials and Human Cell Lines

UT946 (female) cell line was derived from a primary OCT tumor with evidence of anaplastic/poorly differentiated transformation. Snap-frozen and fresh tissue samples were collected under IRB Protocols STU 072017-103 (PI – McFadden) and STU 102010-051 (PI – Lewis). Fresh tissue was minced and placed in 5 mL DMEM (Sigma D6429) containing 200 μL Enzyme H, 100 μL Enzyme R, and 25 μL Enzyme A that were prepared according to the manufacturer’s instruction (Miltenyi 130-095-929) and incubated for approximately 30 minutes at 37°C with agitation every 5-10 minutes. The resulting cell suspension was filtered through a 40 μm nylon mesh filter, pelleted, and resuspended in DMEM supplemented with 10% FBS (Sigma F0926), 2 mM L-glutamine (Sigma G7513), 1% penicillin/streptomycin (Sigma P4333), and 50 μg/mL uridine (Sigma U3003). NCI-237 and UT354 cell lines were generated as previously described in Frank *et al*. 2023^27^. TPC-1 (female) was obtained from Dr. Sareh Parangi (Massachusetts General Hospital, Boston, MA, USA). HEK293T/17 were purchased from ATCC (CRL-11268). HCT-116 (male) was obtained from Dr. Deepak Nijhawan (University of Texas Southwestern Medical Center, TX, USA).

Cell lines were maintained at 37°C with 5% CO2 and cultured in DMEM (Sigma D6429) supplemented with 10% FBS, 2 mM L-glutamine and 1% penicillin/streptomycin unless stated otherwise. For metabolic assays (Figure 1), human plasma-like media (HPLM) was used. Cell lines were adapted to culture in HPLM supplemented with 2% FBS, 1% penicillin/streptomycin, and 1× insulin–transferrin–selenium for at least 5 passages. HPLM pool stocks were prepared and stored as previously described in ^27^. Prepared HPLM was used within 3 to 4 days and supplements were added daily before use. All cell lines were subjected to short-tandem repeat (STR) profiling performed by the Eugene McDermott Center for Human Growth and Development Sequencing Core. Cell lines were periodically tested for mycoplasma contamination using a PCR-based assay. Before conducting cell culture experiments, cell lines were passed at least once after thawing, and experiments were performed within 10 to 25 passages of being thawed.

### Respiration analysis

Mitochondrial respiration measurements were performed using an Agilent Seahorse XFe96 Analyzer according to manufacturer’s instructions for Mito Stress kit. Briefly, cells were plated at 20,000 cells/well in 80 μL of culture media and incubated overnight. The following day, cells were rinsed twice using Seahorse assay medium (DMEM (Sigma D5030) with 10 mmol/L glucose, 2 mmol/L L-glutamine, 1 mmol/L sodium pyruvate, and 1% penicillin/streptomycin), and 150 μL assay medium was added after the second wash. Oxygen Consumption Rate (OCR) and Extracellular Acidification Rate (ECAR) were measured after equilibration in a CO2-free incubator for 45 minutes following the sequential addition of oligomycin (1.5 µM), CCCP (3 µM), Rotenone (0.5 µM) and antimycin A (0.5 µM). For normalization Hoech 33342 10ug/mL was added directly into the wells and incubated for 15 minutes. Live cells were counted using the Celígo™ 4 channel image cytometer (Nexcelom Bioscience).

### Cell proliferation assays

For 3 days assays with multiple cell lines, cells were plated in 6 well plates (Corning) in 2 mL of HPLM media at the following densities: NCI-237 (30,000 cells/well); UT946 (30,000 cells/well); TPC-1 (30,000 cells/well); UT354 (100,000 cells/well); and allowed to adhere overnight. The following day, cells were washed once with 2 mL PBS (Sigma D8537), and 4-8 mL of HPLM growth media was added to the respective wells. A separate “Pool 9 stock” solution that lacked pyruvate was prepared and used to prepare HPLM without pyruvate before pH adjustment and sterile filtration.

For 3- and 5-day proliferation assays, cells of the same type were seeded in duplicate 12-well plates (Corning) at a density of 25,000 cells per well in 1 mL of media. This included TPC-1-derived cells and cybrids, as well as UT946-derived cells and cybrids. The following day (designated as day 0), one plate was washed once with PBS (Sigma, D8537), trypsinized, and counted to establish baseline cell numbers. The remaining plate was also washed with PBS and incubated for either 3 or 5 additional days in 3 mL of the corresponding media. Media lacking specific nutrients (pyruvate, uridine, or glucose) was prepared using glucose-free, pyruvate-free DMEM (Thermo Fisher #11966) supplemented with 10% dialyzed FBS, 2 mM L-glutamine and 1% penicillin/streptomycin as a base. Complete media was supplemented with 1 mM sodium pyruvate (Sigma Aldrich #P2256-10MG), 50 µg/mL uridine (Sigma Aldrich #U3003-50G), and 5 mM glucose (Sigma Aldrich #G7021). To generate nutrient-depleted conditions, the corresponding component (pyruvate or uridine) was simply omitted. For glucose-deprivation experiments, glucose was replaced with 5 mM galactose (Sigma Aldrich #G5388-100G). Conditions were assayed in triplicate.

### Viability assays

Cells were plated in 6 well plates (Corning) in 2 mL of HPLM media at 100,000 cells/well and allowed to adhere overnight. The following day, cells were washed once with 2 mL PBS and 8 mL of HPLM growth media was added to the respective wells. Growth media lacking glucose was prepared by supplementing the media with 5 mM galactose or 5 mM glucose from sterile 500 mM stocks. Cells were cultured for 48 hours before all cells (floating and adherent) were collected by trypsinization and pelleting at 500x *g* for 5 minutes. Cells were resuspended in 500 μL PBS + 0.5 μg/mL propidium iodide (Life Technologies P1304MP) and stained in the dark at room temperature for 10 min before data acquisition and data analysis on a Guava easyCyteTM flow cytometer (EMD Millipore). Conditions were assayed in triplicate and experiments were performed at least twice.

### 13C glucose labelling

Cells were plated in 6 well plates at the following densities: NCI-237 (300,000 cells/well), UT946 (300,000 cells/well), TPC-1 (400,000 cells/well) and UT354 (800,000 cells/well), in 3 mL media and allowed to adhere overnight. The following day, HPLM was prepared and supplemented with 5 mM [U-^13^C_6_] glucose before sterile filtration. Following incubation at 37°C and 5% CO2, cells were collected as described in^27^. Briefly, cells were extracted in methanol:water (80:20) and vacuum-dried before derivatization. Derivatized samples were transferred to Agilent GC-MS autosampler vials, and data acquisition was performed using an Agilent G2579A MSD coupled to an Agilent 6890 gas chromatogram. Data were analyzed using Agilent MSD Chemstation software and a MatLab script for area-under-the-curve analysis, total ion count (TIC) determination, and natural isotopomer abundance correction. Conditions for all GC-MS experiments were assayed in triplicate.

### WES analysis – LOH analysis

For the re-analysis of published OCT data^19,70^, raw reads are trimmed with Trim Galore (https://www.bioinformatics.babraham.ac.uk/projects/trim_galore/) to remove adapter sequences and low-quality bases, then aligned to the human reference genome (hg38) using Burrows-Wheeler Aligner (BWA, v0.7.17)^71^. PCR duplicates are removed with Picard (2.21.3) (https://broadinstitute.github.io/picard), and base-quality scores are recalibrated with Genome Analysis Toolkit (GATK, 4.1.4.0)^72,73^. Variant calling follows GATK best-practice recommendations, and resulting VCFs are filtered (QD < 2, FS > 60, MQ < 40, DP < 10, GQ < 20) and annotated with RefSeq, dbSNP, COSMIC, ClinVar, gnomAD, and CADD using the custom Annomen pipeline (https://github.com/jiwoongbio/Annomen). Genome-wide loss of heterozygosity and runs of homozygosity are inferred from variant-allele-frequency profiles with AutoMap^74^ and validated with PLINK (--homozyg)^75^. Broad LOH regions are visualized through per-chromosome variant-allele-frequency plots generated by a custom R script. Previously published OCT datasets were obtained from Sequence Read Archive (SRA #SRP136351 for Ganly *et al.* 2018) and from the database of Genotypes and Phenotypes (dbGaP:phs001580.v1.p1 for Gopal *et al.* 2018).

### Ploidy

For assessing ploidy in UT946 and HCT116, 500,000 cells were stained using Guava cell cycle Reagent (#4500-0220) following the manufacturer’s instructions. Briefly, cells were centrifuge and washed 1X with PBS. Then cells were added to 70% Ethanol to a concentration of 5x10^5^ cells/mL for fixation and incubated for 3h at −20°C. An aliquot of fixed cells was then centrifuged and washed with 1x PBS. Finally, cells were resuspended in 200 µL of Guava cell cycle Reagent and incubated for 30 min prior to data acquisition and data analysis on the Guava easyCyteTM flow cytometer (EMD Millipore).

### Cytoplasmic Hybrid (Cybrid) Generation

Cybrids were generated using a modified version of the protocol described in [7]. The day prior to cybrid generation, ultracentrifugation tubes were prepared with a Percoll (Sigma, P1644-25ML):media gradient. The media consisted of DMEM (Sigma, D6429) supplemented with 10% FBS, 2 mM L-glutamine, 1% penicillin/streptomycin, and 50 μg/mL uridine. The gradient was layered as follows: 10 mL of 50% Percoll, followed by 4 mL each of 35%, 30%, 26%, and 22% Percoll. Tubes were equilibrated overnight at 37 °C and 5% CO₂. The next day, one 150 mm plate containing 5–20 million donor cells was trypsinized, collected, and counted. Three million cells were set aside as positive and negative controls. The remaining cells were centrifuged at 200 × g for 5 minutes, resuspended in 4 mL of 12.5% Percoll:media solution containing 10 µg/mL cytochalasin B (Cayman, 11328), 100 nM MitoTracker Green (Cell Signaling, #9074P), and 10 µg/mL Hoechst 33342. The suspension was gently layered on top of the prepared gradient. Tubes were balanced and centrifuged at 25,000 rpm for 60 minutes at 37 °C with minimal braking. During centrifugation, one million of the reserved cells were stained with either Hoechst or MitoTracker (single-color controls), and one million remained unstained as a negative control. After centrifugation, the cytoplast-enriched band appearing between gradient layers (∼1 mL) was carefully collected and washed in 49 mL of DMEM (Sigma, D6429). Cytoplasts and controls were centrifuged at 650 × g for 10 minutes, resuspended in FACS buffer (HBSS, Gibco 14175079) containing 2% FBS, 1% penicillin/streptomycin, and 0.5 mM EDTA (Invitrogen, 15575020), and adjusted to ∼5 million cells/mL. Cytoplasts were sorted using a BD FACSMelody™ Cell Sorter by selecting MitoTracker-positive (FITC) and Hoechst-negative (BV421) populations. In parallel, a 100 mm plate of recipient ρ⁰ cells was harvested and counted. Approximately one million ρ⁰ cells were set aside as negative controls. The remaining cells were stained in media containing 50 nM SYTO62 (Invitrogen, #811344) for 1 hour at 37 °C. Cells were then washed, resuspended in 1–2 mL of DMEM, and counted. Following cytoplast sorting, equal numbers of cytoplasts and SYTO62-labeled ρ⁰ cells were mixed in 2 mL of serum-free DMEM and incubated at room temperature for 10 minutes to facilitate membrane contact. A 200 µL aliquot of the mixture was reserved as a non-fused control. Both the fusion mixture and non-fused control were centrifuged at 650 × g for 10 minutes. The non-fused pellet was resuspended in 300 µL of FACS buffer. For the fusion reaction, the supernatant was carefully aspirated from the pellet to avoid disruption. A total of 100 µL of PEG solution was added directly to the pellet, and the cells were gently homogenized. After 1 minute of incubation, pre-warmed serum-free DMEM was gradually added: 100 µL in the first minute, 200 µL in the second, and up to 500 µL total over several minutes. The cell suspension was then incubated at 37 °C for 15 minutes in a water bath, followed by centrifugation at 650 × g for 10 minutes. The resulting fused cell pellet was resuspended in 500 µL of FACS buffer and sorted on the BD FACSMelody™ Cell Sorter. Successfully fused cybrids were identified as double-positive for MitoTracker Green (FITC) and SYTO62 (PE-Cy5). Sorted cells were centrifuged again at 650 × g for 10 minutes, resuspended in DMEM (Sigma, D6429) supplemented with 10% FBS, 2 mM L-glutamine, 1% penicillin/streptomycin, and 50 μg/mL uridine, and plated into a 96-well plate. Media was refreshed the following morning to allow cybrid attachment. Once the cells were established and expanded over several passages, cybrids were transitioned to uridine-free media to eliminate any residual ρ⁰ cell contamination. Uridine-free media was prepared using DMEM (Sigma, D6429) supplemented with 10% dialyzed FBS, 2 mM L-glutamine, and 1% penicillin/streptomycin. Cybrids were cultured in this selective media for 4–6 passages, after which mitochondrial and nuclear DNA content was assessed. All cybrids were subjected to short-tandem repeat profiling performed by the Eugene McDermott Center for Human Growth and Development Sequencing Core to validate the nuclear entity of the new cybrid line.

### ρ⁰ Cell Generation

Cells were cultured in complete medium consisting of DMEM (Sigma, D6429) supplemented with 10% FBS, 2 mM L-glutamine, 1% penicillin/streptomycin, and 50 μg/mL uridine. To deplete mitochondrial DNA (mtDNA), cells were treated with 10 μM 2’,3’-dideoxycytidine (ddC) (MedChem, HY-17392) and passaged every two days. mtDNA depletion was monitored by quantitative PCR (qPCR) and considered successful when reduced by more than 500-fold relative to wild-type cells. Since mtDNA levels slowly recover after ddC removal, ρ⁰ cells were used shortly after drug withdrawal, and mtDNA abundance was routinely quantified during cybrid generation.

### DNA Extraction

Genomic DNA was isolated from tissue culture samples and snap-frozen tissue samples by resuspension in approximately 500 mL genomic DNA extraction buffer (10 mM Tris-HCl pH 8.0, 25 mM EDTA pH 8.0, 100 mM NaCl, and 1% SDS) supplemented with 0.5 mg/mL proteinase K (Zymo, D3001-2-20) and incubated overnight at 55 °C. DNA was subsequently purified by phenol–chloroform extraction followed by ethanol precipitation as previously described^76^. The resulting DNA was resuspended in nuclease-free water or TE buffer (1 mM EDTA, pH 8.0; 10 mM Tris, pH 8.0) and stored at −20 °C for downstream applications. For genomic DNA isolation from FFPE samples, consecutive 10 mm sections were mounted on clean slides. Tumor and normal thyroid margins were demarcated by a trained pathologist (J. Bishop). The respective tumor and normal thyroid sections were gently scraped from slides using a clean scalpel and transferred to clean tubes. Samples were processed using the QIAamp DNA FFPE Tissue Kit (Qiagen 56404) according to the manufacturer’s instructions.

### RNA Extraction and cDNA Synthesis

Total RNA was extracted using TRIzol reagent (Life Technologies, 15596-018) following the manufacturer’s protocol. One microgram of RNA was reverse-transcribed into cDNA using the High Capacity cDNA Reverse Transcription Kit (Applied Biosystems, #4368813) according to the manufacturer’s instructions. Synthesized cDNA was stored at −20 °C for future analyses.

### Quantitative PCR (qPCR)

qPCR was performed in triplicate using the GoTaq qPCR Master Mix (Promega, A6001) in a total reaction volume of 10 μL, containing 3 ng of DNA or cDNA. For mtDNA quantification, the nuclear-encoded gene ACTB was used as a reference for nuclear DNA. Mitochondrial genes ND1 and COX3 served as proxies for mtDNA content. NDUFA6 was used for gene expression analysis. See Supplementary Table 4 for primers

### mtDNA Sequencing Validation

To validate mtDNA in cybrids, long-range PCR was performed using total DNA as template and KAPA HiFi HotStart DNA Polymerase (Roche, 07958927001). Amplified mtDNA fragments were submitted to Plasmidsaurus for linear PCR sequencing using Oxford Nanopore Technology, with custom analysis and variant annotation (Supplementary Table 4)

### PCR amplification and Sanger Sequencing

The NDUFS1 V228A mutation was validated by PCR amplification of exon 8 using total DNA as a template. PCR products were subjected to Sanger sequencing with the forward primer used as the sequencing primer. To confirm the expression of the correct coding sequences (wild-type vs. mutant) for the seven missense mutations expressed from PGK-driven plasmids in TPC KO cells, total DNA was amplified by PCR using the corresponding primer pairs (Supplementary Table 4). All PCR reactions were performed with KAPA HiFi HotStart DNA Polymerase (Roche, 07958927001). Mutations were verified by Sanger sequencing using at least the forward primer; when necessary, additional primers were employed to confirm the presence or absence of the mutation of interest.

### Lentivirus production

Lentiviruses were produced by co-transfecting HEK 293T/17 cells with the lentiviral expression plasmid of interest along with the packaging plasmids psPAX2 and pMD2.G at a 5:3:2 mass ratio (vector:psPAX2:pMD2.G), using TransIT-LT1 transfection reagent (Mirus, MIR 2300). Viral supernatants were collected at 48 and 72 hours post-transfection, pooled, and filtered through 0.45 μm PES filters. Supernatants were either used immediately or aliquoted and stored at –80 °C for future use.

### Generation of CRISPR-Cas9–edited cells

To knock out the desired genes (NDUFS1, NDUFV1, NDUFAF6, NDUFS8, NDUFA6, NUBPL, and TMEM126B), three sgRNAs per gene were designed using CrisPick (Broad Institute) and/or based on the Brunello library and cloned into the LentiCRISPR-v2 backbone. Briefly, LentiCRISPR-v2 was linearized with BsmBI-v2 (New England Biolabs, #R0739L) according to the manufacturer’s instructions, and the linearized plasmid was gel-purified using the Macherey-Nagel Gel Extraction Kit (#740609.50S). In parallel, sgRNA oligonucleotides were annealed and phosphorylated with T4 Polynucleotide Kinase (New England Biolabs, #M0201S) and ligated into the linearized vector using T4 DNA Ligase (New England Biolabs, #M0202S). Recombinant plasmids were transformed into Stbl3 competent cells and grown under ampicillin selection (100 μg/mL). Plasmid DNA was isolated using the Promega Miniprep Kit (#A1222). Lentiviruses encoding sgRNAs were produced as described above. TPC-1 cells were seeded at 4 × 10⁵ cells per well in 6-well plates, and lentiviral supernatants were added at a 1:1 ratio with fresh medium at the time of plating. After overnight incubation, the medium was replaced, and 48 hours post-transduction, cells were transferred to selection medium containing 2 μg/mL puromycin (Sigma, P8833). Selection was maintained for 3–5 days to generate pooled CRISPR–Cas9–edited populations. Cell survival following puromycin selection was typically 80–100%, as estimated visually. Protein depletion was verified by Western blotting, and the most efficient sgRNA for each gene (listed in Supplementary Table 4) was selected for downstream experiments. When population-level results were inconclusive, single-cell clones were isolated from the total population. Briefly, 500, 1,000, or 2,000 TPC-1 cells were seeded in independent 150 cm² plates in triplicate. After approximately two weeks, individual colonies were manually picked, expanded, and validated by Western blotting. Verified clones were subsequently used for re-expression of either the wild-type or mutant form of the gene of interest.

### Cloning strategy for stable expression of complex I genes

To stably express wild-type (WT) or mutant forms of human Complex I genes (NDUFS1, NDUFV1, NDUFAF6, NDUFS8, NDUFA6, NUBPL, TMEM126B) in CRISPR-Cas9–edited cells, point mutations were introduced by PCR amplification. These included silent mutations in the protospacer adjacent motif (PAM) sequences to prevent re-editing, as well as the desired missense mutations. Reference human cDNA templates were obtained from the McDermott Center for Human Growth and Development (Ultimate™ ORF Lite human cDNA collection, Life Technologies) for NDUFS1, NDUFV1, NDUFA6, NDUFS8, NUBPL and used as PCR templates. For NDUFAF6 and TMEM126B, cDNA templates were obtained from GeneScript (Clone ID: OHu19552D and OHu29970D respectively). Gibson assembly was used (New England biolabs #E2621S) to insert amplified cDNA fragments into a PGK-IRES-Blast lentiviral vector previously digested with BamHI and XBaI (New England biolabs #R3136S and #R0145S). Amplified cDNA fragments were inserted into a PGK-IRES-Blast lentiviral vector previously digested with BamHI and XBaI (New England Biolabs, #R3136S and #R0145S) using Gibson Assembly (New England Biolabs, #E2621S), generating the final expression constructs. For re-expression of WT or mutant constructs in TPC-1 knockout cells, lentiviral transduction was performed as described for CRISPR–Cas9–mediated gene editing. Forty-eight hours post-transduction, cells were plated directly into medium containing 5 μg/mL blasticidin (InvivoGen, ant-bl-1) and maintained under selection for 7–10 days to generate pooled re-expressing cell lines. Cell survival following blasticidin selection was typically 80–100%, as estimated visually.

### Repair of NDUFS1^V228A^ using IDT Alt-R™ CRISPR-Cas9 system

Repair of the NDUFS1^V228A^ point mutation directly in UT946 cells was achieved via homology-directed repair (HDR) using the Alt-R™ CRISPR-Cas9 System (IDT) and single-stranded donor oligonucleotides (ssODNs). HDR donor templates flanking the V228A mutation in exon 8 of *NDUFS1* were designed using the Alt-R HDR Design Tool (IDT) ^77^. The protocol involved co-delivery of the HDR donor ssODN and a CRISPR-Cas9 ribonucleoprotein (RNP) complex via electroporation using the 4D-Nucleofector™ System (Lonza). For nucleofection, 1.5 million UT946 cells were washed with 1× PBS and resuspended in 71.4 µL of Buffer SE (Lonza). The cell suspension was combined with the RNP complex, and HDR donor template, transferred to a well of a Nucleocuvette™ Plate (Lonza), and electroporated using program EO-100. After nucleofection, cells were rested for 10 minutes at room temperature and then transferred to 100-mm dishes containing pre-warmed complete media. Cells were incubated at 37 °C and 5% CO₂, and media was replaced the following morning. After several passages and complete recovery, selection for successfully repaired cells was performed by culturing cells for 24 hours in glucose-free, galactose-containing medium (DMEM D5030 supplemented with 10% dialyzed FBS, 1 mM sodium pyruvate, 5 mM galactose, 2 mM L-glutamine, 1% penicillin/streptomycin, and 50 μg/mL uridine). Remaining viable clones were expanded in regular media and validated by Sanger sequencing.

### Immunoblotting

Cells were rinsed with ice-cold PBS and lysed in buffer containing 10 mM HEPES (pH 7.4), 150 mM NaCl, 2 mM MgCl₂, and 1% SDS, supplemented with 0.08 units of Universal Nuclease (Thermo Fisher, #88702). Lysates were incubated for 10 minutes at room temperature, followed by boiling at 96 °C for 10 minutes. Protein content was measured with BCA (Bicinchoninic Acid) Protein Assay (Thermo Scientific™ 23222), protein was denatured by the addition of sample buffer, boiled for 5 min, resolved by SDS–PAGE and analyzed by immunoblotting. Western blot analyses were performed according to standard procedures. Antibodies were visualized by using Odyssey Infrared Imaging System (Application software version 3.0.30) LI-COR Biosciences. Antibodies from Cell Signaling Technology were used for detection of β-Actin (#4970) dilution 1:5000 and P-AMPK (#2535) dilution 1:1000. Antibodies from Proteintech were used to detect NDUFV1 (#11238-1-AP) dilution 1:1000, Vinculin (#66305-1-Ig) dilution 1:5000, dilution 1:1000 and NDUFA6 (#15445-1-AP) dilution 1:500. Antibodies from Abcam were used to detect NDUFS1 (#ab169540) dilution 1:1000, NDUFS3 (#ab110246) dilution 1:1000, NDUFS8 (#sc-515527) dilution 1:500 and NUPBL (#EPR11833) dilution 1:1000. The antibody from Millipore Sigma was used to detect NDUFAF6 (#HPA050545).

## Acknowledgments

We thank Dr. Spencer D. Shelton and Dr. Prashant Misra for sharing the protocol of cytoplasmic hybrids generation. We thank Cheryl Lewis and the Simmons Comprehensive Cancer Center Tissue Management Shared Resource for collecting and providing tumor samples. We thank members of the McFadden lab for helpful discussions and review of the manuscript.

## Author contributions

Conceptualization: CCA, AF, DGM

Methodology: CCA, AF, DGM

Investigation: CCA, AF, KMu, JK, CL, KMa, JAB

Visualization: CCA

Supervision: DGM, YX

Writing—original draft: AF, CCA

Writing—review & editing: CCA, AF, DGM

## Data and materials availability

WES data from UT946 tumor and cell line will be deposited in an approved repository prior to publication

