## Supplementary Materials for "Loss of heterozygosity exposes germline mutations in complex I and drives Warburg metabolism in oncocytic carcinoma of the thyroid"

Celia de la Calle Arregui *et al.*

*

**This PDF file includes:**

Figs. S1 to S4
Table S6

#### SUPPLEMENTARY FIGURE 1

**A**

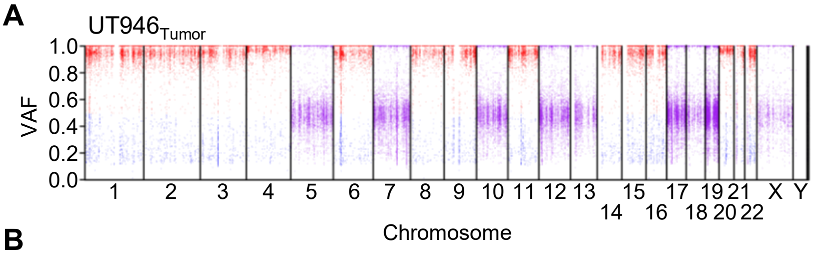


**B**

$UT946_{Tumor}$

| Gene | Mutation | Protein alteration | VAF | Clinical Variant significance | |
|---|---|---|---|---|---|
| BRCA1 | chr17:43104249T>C | p.Tyr105Cys | 0.466 | Flagged | Benign |
| DNMT3A | chr2:25300227T>G | p.Glu30Ala | 0.990 | Flagged | Benign. Likely benign |

$UT946_{Cell\ Line}$

| Gene | Mutation | Protein alteration | VAF | Clinical Variant significance | |
|---|---|---|---|---|---|
| DNMT3A | chr2:25300227T>G | p.Glu30Ala | 1.000 | Flagged | Benign/Likely benign |
| BRCA1 | chr17:43104249T>C | p.Tyr105Cys | 0.983 | Flagged | Benign |
| PIK3CA | chr3:179199088G>A | p.Arg88Gln | 0.451 | Flagged | Likely pathogenic |
| CDKN2A | chr9:21971156G>A | p.Ala68Val | 0.565 | Flagged | Conflicting interpretations of pathogenicity |
| PTEN | chr10:87957915C>T | p.Arg233* | 0.960 | Flagged | Pathogenic |
| TP53 | chr17:7674220C>T | p.Arg248Gln | 0.950 | Flagged | Pathogenic Likely pathogenic |
| NF2 | chr22:29661313C>T | p.Arg262* | 0.992 | Flagged | Pathogenic |

**C**

$UT946_{Tumor}$

| Gene | Mutation | Protein alteration | VAF | ClinVar significance | |
|---|---|---|---|---|---|
| MT-ND5 | ChrM:12397A>G | p.Thr21Ala | 0.999 | Flagged | Benign |
| MT-TRNQ | chrM:4336T>C | non-CDS | 1.000 | Flagged | Conflicting interpretations of pathogenicity |

$UT946_{Cell\ Line}$

| Gene | Mutation | Protein alteration | VAF | ClinVar significance | |
|---|---|---|---|---|---|
| MT-ND5 | ChrM:12397A>G | p.Thr21Ala | 1.000 | Flagged | Benign |
| MT-TRNQ | chrM:4336T>C | non-CDS | 1.000 | Flagged | Conflicting interpretations of pathogenicity |
| MT-CYTB | ChrM:15884G>A | p.Ala380Thr | 0.432 | Flagged | Benign |
| MT-ND1 | ChrM:4210G>A | p.Ala299Thr | 0.233 | | |
| MT-COX3 | ChrM:9804G>A | p.Ala200Thr | 0.180 | Flagged | Conflicting interpretations of pathogenicity |

**Supplementary Figure 1. A,** Loss of heterozygosity plot showing variant allele fraction (VAF) of germline variants for UT946 Tumor. Variants with VAF >0.5 are in red; variants with VAF <0.5 are in blue. **B,** Table showing selected somatic mutations in nuclear-encoded genes in UT946 tumor (up) and cell line (down). **C,** Table showing selected somatic mutations in mitochondrial-encoded genes in UT946 tumor (up) and cell line (down).

### SUPPLEMENTARY FIGURE 2

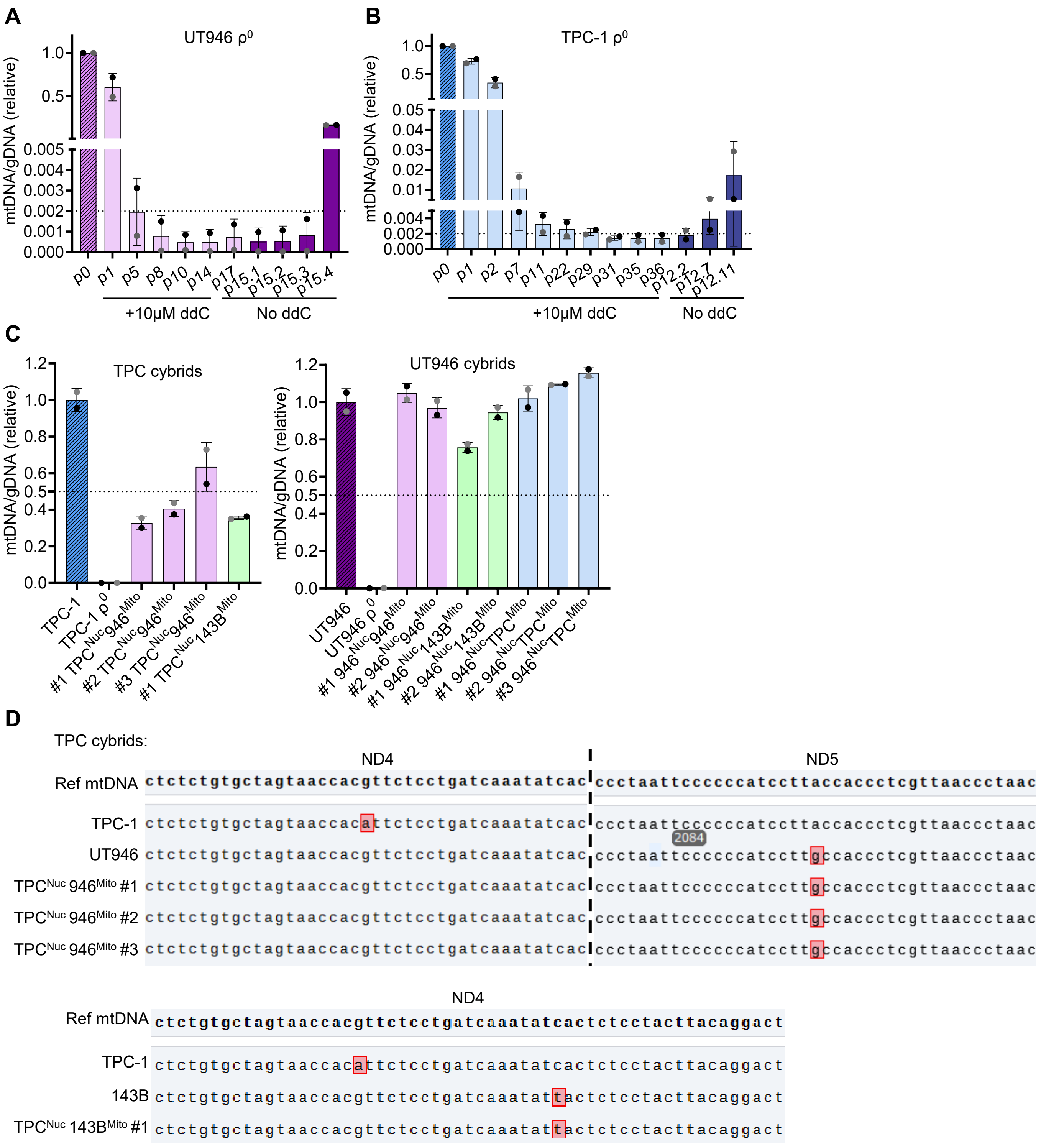

#### SUPPLEMENTARY FIGURE 2

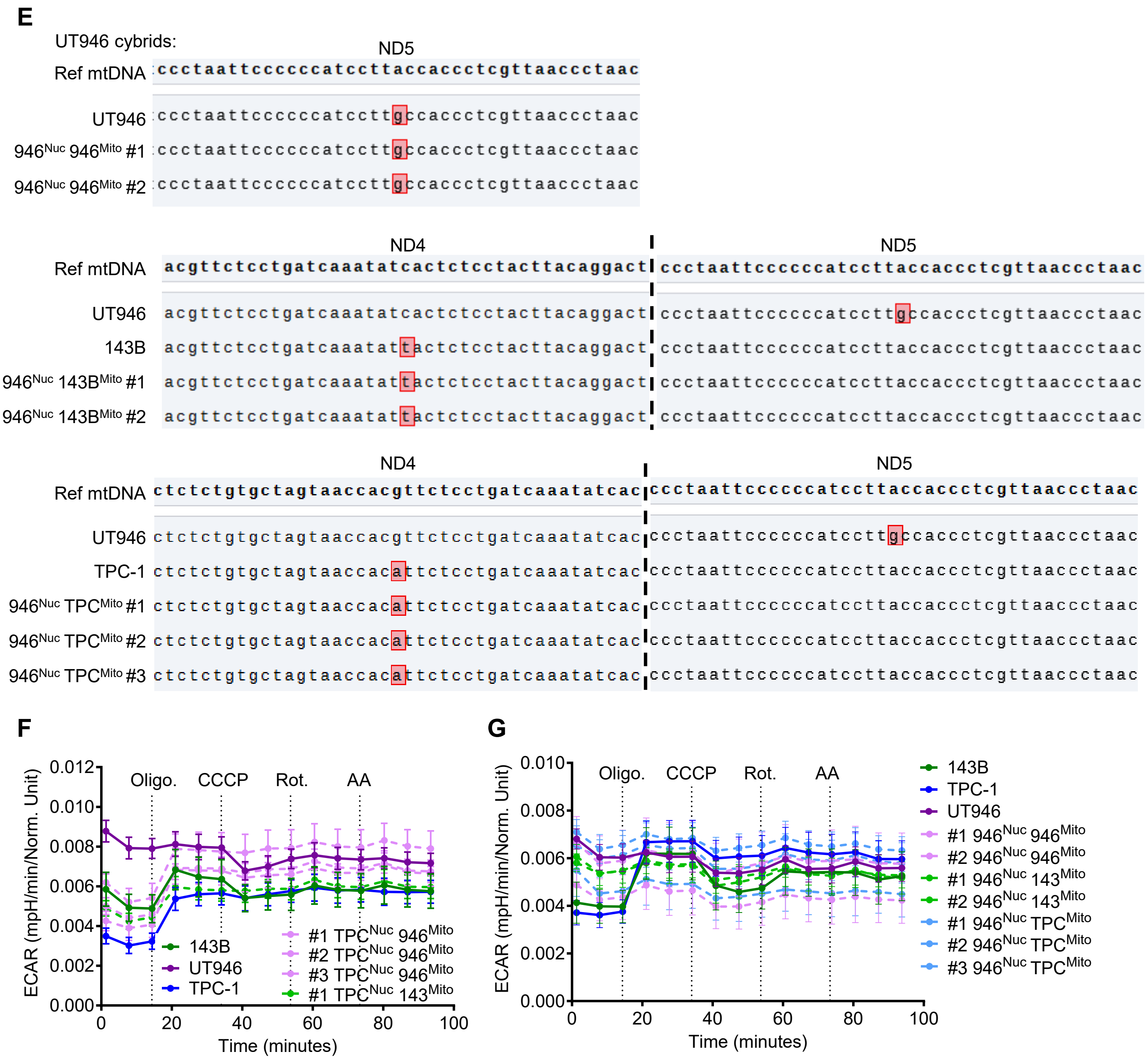


**Supplementary Figure 2. A-C,** Relative mtDNA levels assessed by measuring the expression levels of *ND1* (gray dots) and *COX3* (black dots) genes normalized to the expression levels of nuclear DNA *ACTB* gene. Total DNA from indicated cell lines and cybrids was used as the template. Cell passages indicated when necessary (p0, p1 etc.) **D-E,** Oxford nanopore long-read sequencing of PCR-amplified mtDNA fragments from indicated cybrids and WT cell lines. **F-G,** Acidification rate in intact cells and cybrids after the addition of Oligo = oligomycin; CCCP = carbonyl cyanide 3-chlorophenylhydrazone; Rot. = Rotenone and Anti A = antimycin A. n=8-12. Data are plotted as mean ± SD of the indicated number of replicates.

#### SUPPLEMENTARY FIGURE 3

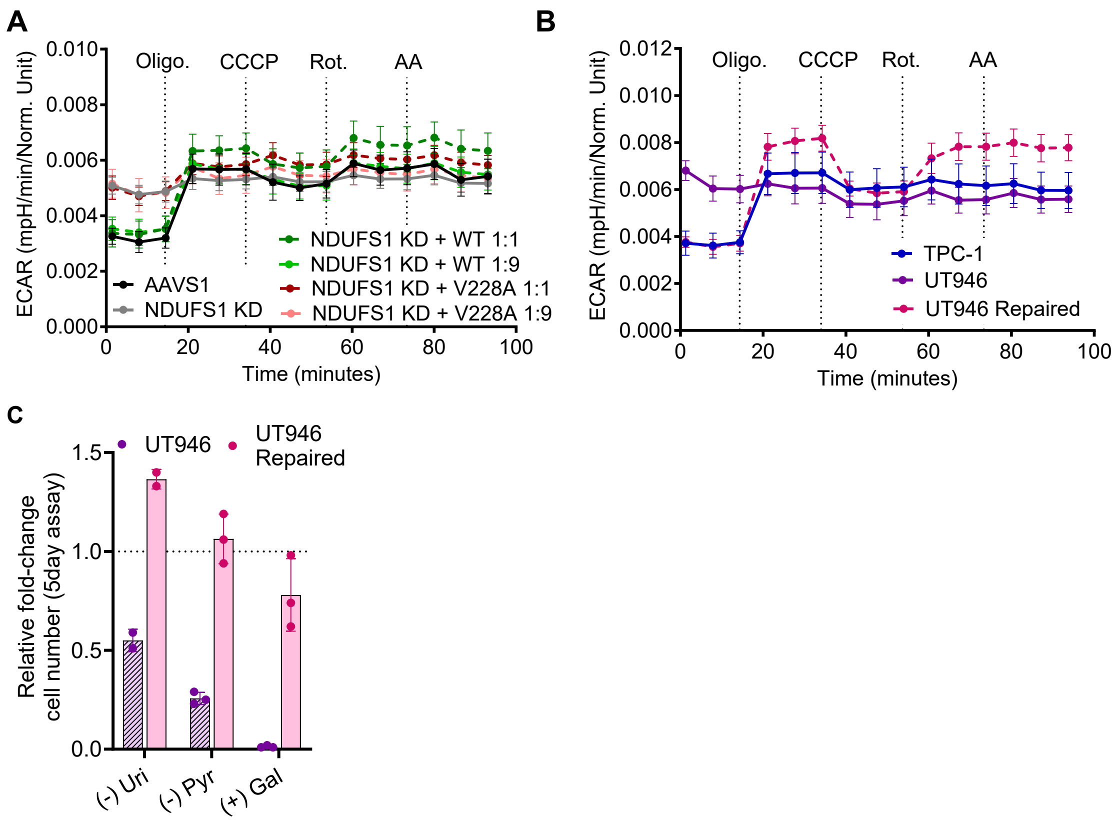


**Supplementary Figure 3. A-B,** Acidification rate in intact indicated cells. 1:1, 1:9 refers to viral dilution upon NDUFS1 re-expression. Oligo = oligomycin; CCCP = carbonyl cyanide 3-chlorophenylhydrazone; Rot. = Rotenone and Anti A = antimycin A. n=8-12. **C,** Fold change in cell number relative to complete media of indicated cell lines in media containing ± 100 µmol/L pyruvate (Pyr.), 50 µg/mL Uridine (Uri.), 5 mM glucose or 5mM galactose (Gal.) n = 3. Data are plotted as mean ± SD of the indicated number of replicates

#### SUPPLEMENTARY FIGURE 4

**A**

| Sample | NL tissue | CI gene | Complex I | Mut class | Protein variation |
|---|---|---|---|---|---|
| UT946 | NL thyroid | NDUFS1 | CORE | missense | p.Val228Ala |
| HCTC-2 | Blood | | | | |
| HCTC-6 | NL thyroid | | | | |
| HCTC-117 | NL thyroid | NDUFA6 | Supernum. | missense | p.His108Tyr |
| HCTC-102 | Blood | | | FS | p.Met104Cysfs*35 |
| HCTC-79 | NL thyroid | NDUFAF7 | Asm. factor | FS | p.Gln384Alafs*7 |
| HCTC-53 | Blood | NDUFAF3 | Asm. factor | FS | p.Gly164Serfs*29 |
| HCTC-84 | Blood | NDUFAF6 | Asm. factor | splicing | |
| SRS3086588 | NL thyroid | | | missense | p.Ala178Pro |
| SRS3086592 | NL thyroid | NUBPL | Asm. factor | missense | p.Asn198Thr |
| SRS3086539 | NL thyroid | | Asm. factor | startcodon | p.? |
| SRS3086540 | NL thyroid | NDUFB6 | Supernum. | nonsense | p.Tyr84* |
| SRS3086620 | NL thyroid | NDUFAF4 | Asm. factor | startcodon | p.? |
| SRS3086621 | NL thyroid | NDUFV2 | CORE | FS | p.His21Argfs*6 |
| SRS3086634 | NL thyroid | NDUFS8 | CORE | missense | p.Trp61Arg |
| SRS3086616 | NL thyroid | NDUFV1 | CORE | missense | p.Leu138Pro |
| HCTC-20 | Blood | TMEM126B | Asm. factor | missense | p.Asp133Asn |
| HCTC-3 | Blood | NDUFV3 | Supernum. | missense | p.Pro4Arg |
| SRS3086618 | NL thyroid | | | missense | p.Arg26Gln |

**B**

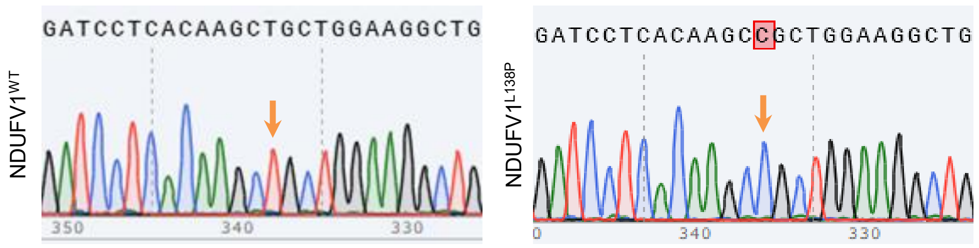


**C**

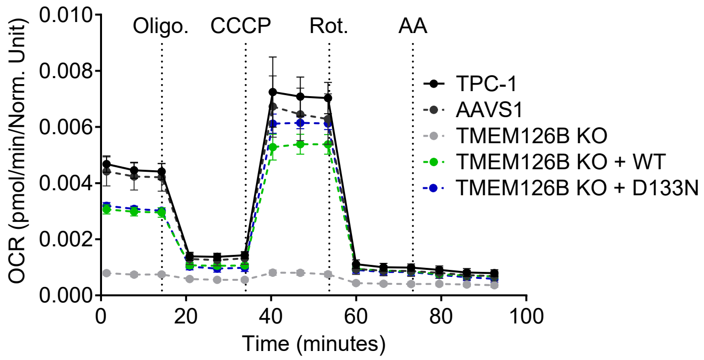


**D**

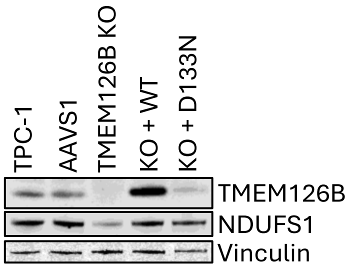


**E**

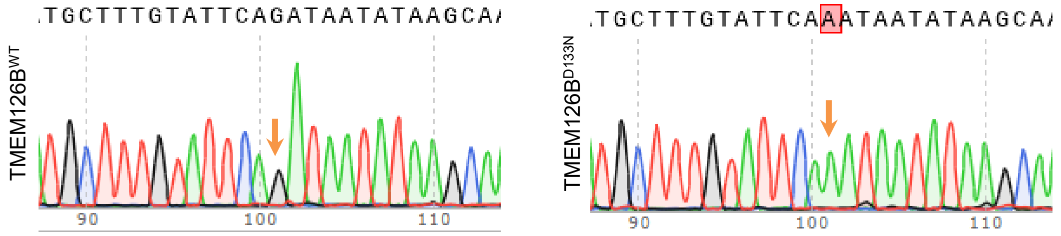

### SUPPLEMENTARY FIGURE 4

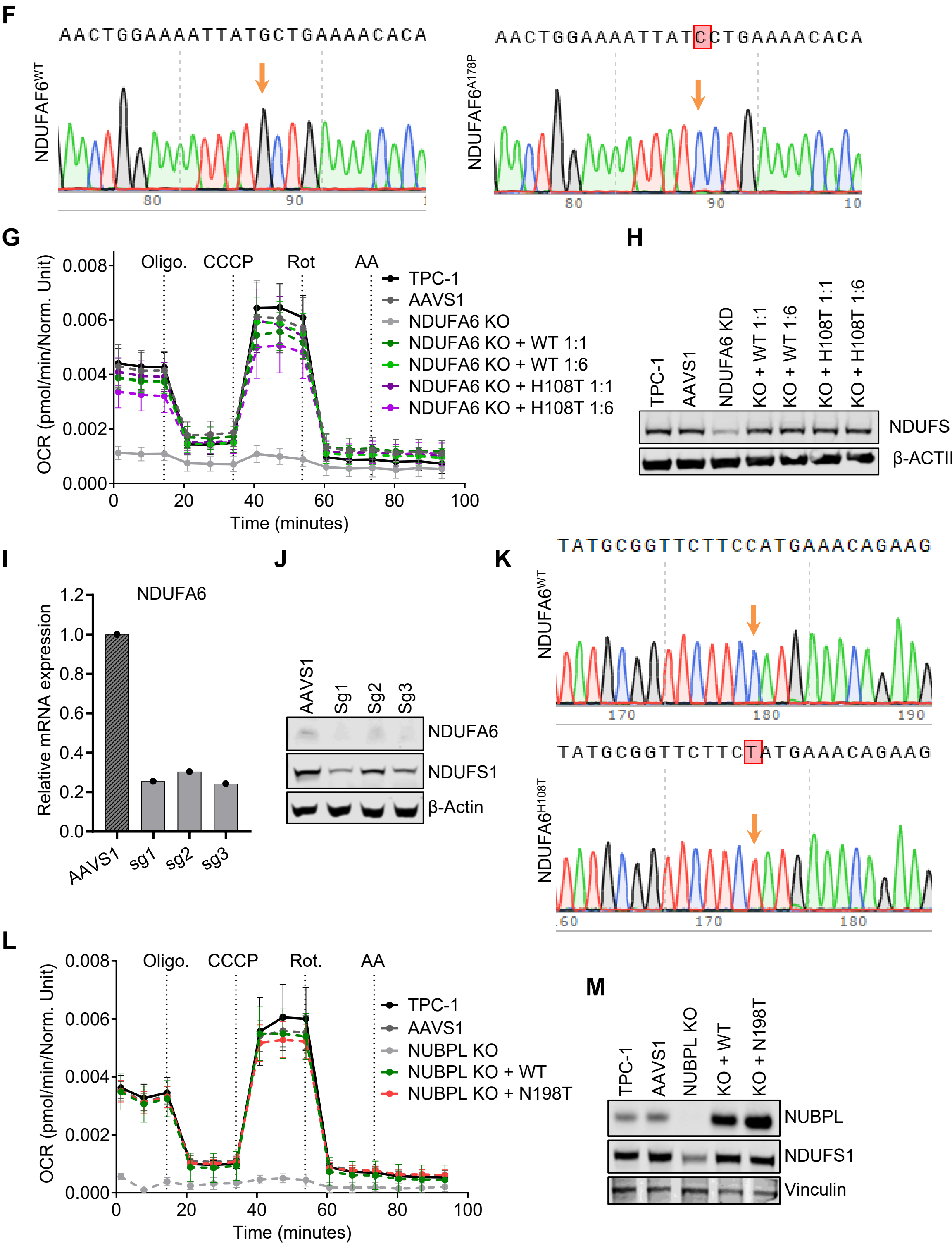

#### SUPPLEMENTARY FIGURE 4

**N**

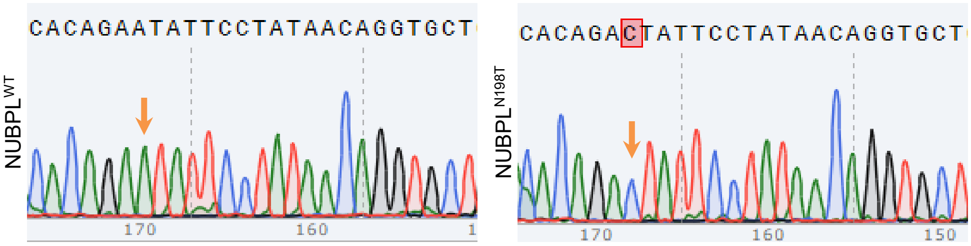


**O**

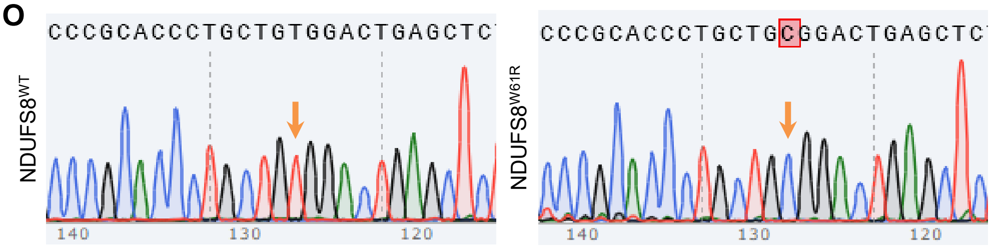


**Supplementary Figure 4. A,** Table showing recessive germline variants that underwent LOH in the tumor targeting Complex I genes encoded in the nucleus. **B,** Sanger sequencing traces of the re-expressed wild type (WT) and mutant form of indicated gene in NDUFV1 TPC-1-edited cells. gDNA use as a template. **C,** Normalized oxygen consumption rate in indicated TPC-1-edited cell lines n=8-12. Oligo = oligomycin; CCCP = carbonyl cyanide 3-chlorophenylhydrazone; Rot. = Rotenone, Anti A = antimycin A. **D,** Protein lysates form TPC-1 edited cells were immunoblotted for indicated proteins **E-F,** Sanger sequencing traces of the re-expressed wild type (WT) and mutant form of indicated gene in TPC-1-edited cells. DNA use as a template. **G,** Normalized oxygen consumption rate in indicated TPC-1-edited cell lines n=8-12. Oligo = oligomycin; CCCP = carbonyl cyanide 3-chlorophenylhydrazone; Rot. = Rotenone, Anti A = antimycin A. **H,** Protein lysates form TPC-1 edited cells were immunoblotted for indicated proteins. **I,** RT-qPCR of cells expressing different single guide RNA (sg) targeting NDUFA6. Expression levels of NDUFA6 relative to β-actin. **J,** Protein lysates form TPC-1 edited cells were immunoblotted for indicated proteins**. K,** Sanger sequencing traces of the re-expressed wild type (WT) and mutant form of NDUFA6 genes in TPC-1-edited cells. DNA use as a template. **L,** Normalized oxygen consumption rate in indicated TPC-1-edited cell lines n=8-12. Oligo = oligomycin; CCCP = carbonyl cyanide 3-chlorophenylhydrazone; Rot. = Rotenone, Anti A = antimycin A. **M,** Protein lysates form TPC-1 edited cells were immunoblotted for indicated proteins. **N-O,** Sanger sequencing traces of the re-expressed wild type (WT) and mutant form of NUBPL and NDUFS8 genes in TPC-1-edited cells. DNA use as a template. Data are plotted as mean ± SD of the indicated number of replicates

#### SUPPLEMENTARY TABLE 4

| Gene | Type | Forward | Reverse |
|---|---|---|---|
| ND3/4 | PCR | TAATAATCTTATATAAACTTCGCCTTAATTTTAATAATCAACACCCT | TGCATGGTTATAAGAGGAAAACCCGGTAATGATGT |
| ND5/6 | PCR | TTTTCCTCTTATAACCATGCACACTACTATAACCACCC | GCCTGCAGGTCGACTCTAGAATGATGTATGCTTTGTTTCTGTTGAGTGT |
| NDUFS1 | PCR | AGACCAAAACAACTGAAACCCA | ACAAGTTTCTGCCTGTGTGT. |
| NDUFV1 | PCR | CAGACGAGGGTGAGCCGGGCAC | GTCCGTCGAGCAGTCCATGACGA |
| NDUFAF6 | PCR | AACTATGGAAAGCTGTTAAAAG | TGTGAAAGGACCTAGCATGC |
| NDUFS8 | PCR | CTGCCTGACCACGCCTAT | TGTTGTACAGCAGCTCCTCA |
| TMEM126B | PCR | CTCAAACTTTCTGTTCAGACGC | GGGGGGAGGGAGAGGGGCGGTTACTCTTCATGTATAGTTTTCTCAAGTGTCT |
| NDUFA6 | PCR | GGAATTCCTCGAGACTAGTTATGGCGGGGAGCGGCGTCC | GGGGGGAGGGAGAGGGGCGGTCATGGATCGTGGCCAACATAGAAC |
| NUBPL | PCR | GGTAATGTCAGCCATTGAGAAATTG | AGTTTCCTTGCACCATCAGC |
| ACTB | qPCR | GGCATCCTCACCCTGAAGTA | AGGTGTGGTGCCAGATTTTC |
| ND1 | qPCR | ACTACAACCCTTCGCTGACG | GCGGTGATGTAGAGGGTGAT |
| COX3 | qPCR | TCCACTCCATAACGCTCCTC | GTGGCCTTGGTATGTGCTTT |
| NDUFA6 | qPCR | CGCCAGCACTTTCGTGAAGC | CTGTTTCATAGAAGAACCGC |
| NDUFS1 | sgRNA | CACCGTAGAGGTTAGGGCACCTACA | AAACTGTAGGTGCCCTAACCTCTAC |
| NDUFV1 | sgRNA | CACCGATCATGGCGTAAGATCTCC | AAACGGAGATCTTACGCCATGATC |
| NDUFAF6 | sgRNA | CACCGTCTGTACGCGCGCATGCGG | AAACCCGCATGCGCGCGTACAGAC |
| NDUFS8 | sgRNA | CACCGCATGAAGTCAGTGACTGACC | AAACGGTCAGTCACTGACTTCATGC |
| TMEM126B | sgRNA | CACCGACTGAGGAAGCGCCCAAGGT | AAACACCTTGGGCGCTTCCTCAGTC |
| NDUFA6 | sgRNA | CACCGACATGAACGAGGCCAAGCGG | AAACCCGCTTGGCCTCGTTCATGTC |
| NUBPL | sgRNA | CACCGCTAGCAGCGAACGATTCGGT | AAACACCGAATCGTTCGCTGCTAGC |
